# Comparative single-cell regulome reveals evolutionary innovations in neural progenitor cells during primate corticogenesis

**DOI:** 10.1101/2023.09.20.558575

**Authors:** Yuting Liu, Xin Luo, Yiming Sun, Kaimin Chen, Ting Hu, Benhui You, Jiahao Xu, Fengyun Zhang, Xiaoyu Meng, Xiang Li, Xiechao He, Cheng Li, Bing Su

## Abstract

The cellular and genetic mechanism underlying the human-specific features of cortex development remains unclear. We generated a cell-type resolved atlas of transcriptome and regulome of the developing macaque and mouse prefrontal cortex, and conducted evolutionary analyses with the published complementary human data. We discovered a primate-specific expansion of two neural progenitor subclasses, glia-committed radial glia (RG) and truncated RG. Specifically, the human neural progenitors show extensive transcriptional rewiring in the growth factor and extracellular matrix pathways. Expression of the human-specific progenitor marker *ITGA2* in the cortex of fetal mouse promotes progenitor proliferation and an increased upper-layer neuron proportion. We demonstrate that these transcriptional divergences are primarily driven by the activity changes of the distal regulatory elements in the genome. Markedly, the chromatin regions with human-gained accessibility enrich the human-fixed sequence changes, as well as sequence polymorphisms associated with intelligence and neuropsychiatric disorders. Our results uncover evolutionary innovations in neural progenitors and gene regulatory mechanism during primate cortex evolution.

## Introduction

The expansion and elaboration of the cortex are thought to underlie the higher cognitive functions emerged during primate evolution (*1, 2*). In particular, the human cortex has acquired several key evolutionary innovations, mainly the increased diversity and enhanced proliferation of progenitor cells, along with the protracted duration of neurogenesis (*3, 4*). These human cortex features likely stem from molecular and cellular rewiring in neural progenitor cells during corticogenesis (*5*). Previous comparative transcriptomic analysis has enabled the identification of these human-specific features (*6–11*). However, further in-depth understanding has been hindered by the limited availability of fetal cortex data from non-human primates (NHPs), and the lack of systematic cross-species comparison of cellular composition and properties of neural progenitors. Moreover, the genetic basis underlying the human cortex innovations remains largely illusive, predominantly due to the limited functional interpretations of the human-specific genomic alterations at cellular resolution during corticogenesis.

To decipher the regulatory mechanism underlying the evolutionary innovations of the human cortex, using single-nucleus multi-omics technology, we simultaneously profiled gene expression and chromatin accessibility of the fetal prefrontal cortex (PFC) of rhesus macaque (*Macaca mulatta*) and mouse (*Mus musculus*) at mid-gestation. Combined with the published human PFC data at matched developmental stage (*12, 13*), our comparative analysis discovered the primate-shared and the human-specific features in the compositions and properties of neural progenitor cells. The cross-species comparison of single-cell regulome demonstrated extensive evolutionary changes in distal regulatory elements during human corticogenesis, and their potential roles in shaping up cognitive traits and brain diseases such as intelligence quotient (IQ) and schizophrenia (SCZ). The presented data offers mechanistic insights into the evolution of human cortex development and pathogenesis of the neuropsychiatric disorders.

## Results

### Single-cell regulatory atlas of the macaque and mouse fetal PFC during corticogenesis

With the use of the single-nucleus approach, we first profiled the gene expression and chromatin accessibility of the prefrontal cortex (PFC) of rhesus macaque (*Macaca mulatta*) at mid-gestation (embryonic days E80-E92) (Fig. 1A). The macaque PFC samples include 6 animals for single-nucleus RNA sequencing (snRNA-seq) and 7 animals for single-cell assay for transposase accessible chromatin with sequencing (scATAC-seq) profiles, and 4 animals for snMultiome sequencing (measuring gene expression and chromatin accessibility from the same nucleus). In addition, to obtain approximate spatial information, using laser microdissection (LMD), we divided the PFC into eight laminae (marginal zone, MZ; outer cortical plate, CPo; inner cortical plate, CPi; subplate, SP, intermediate zone, IZ, outer subventricular zone, OSVZ; inner subventricular zone, ISVZ and ventricular zone, VZ), and generated lamina-bulk RNA-seq data. Also, as cross-data validation for the scATAC-seq, we conducted bulk ATAC-seq of the macaque PFC. In addition, we included our previous macaque bulk Hi-C data (the fetal PFC at the same developmental stage) (*14*) to validate the inferred peak-to-gene links by the snMultiome data.

**Fig. 1.**
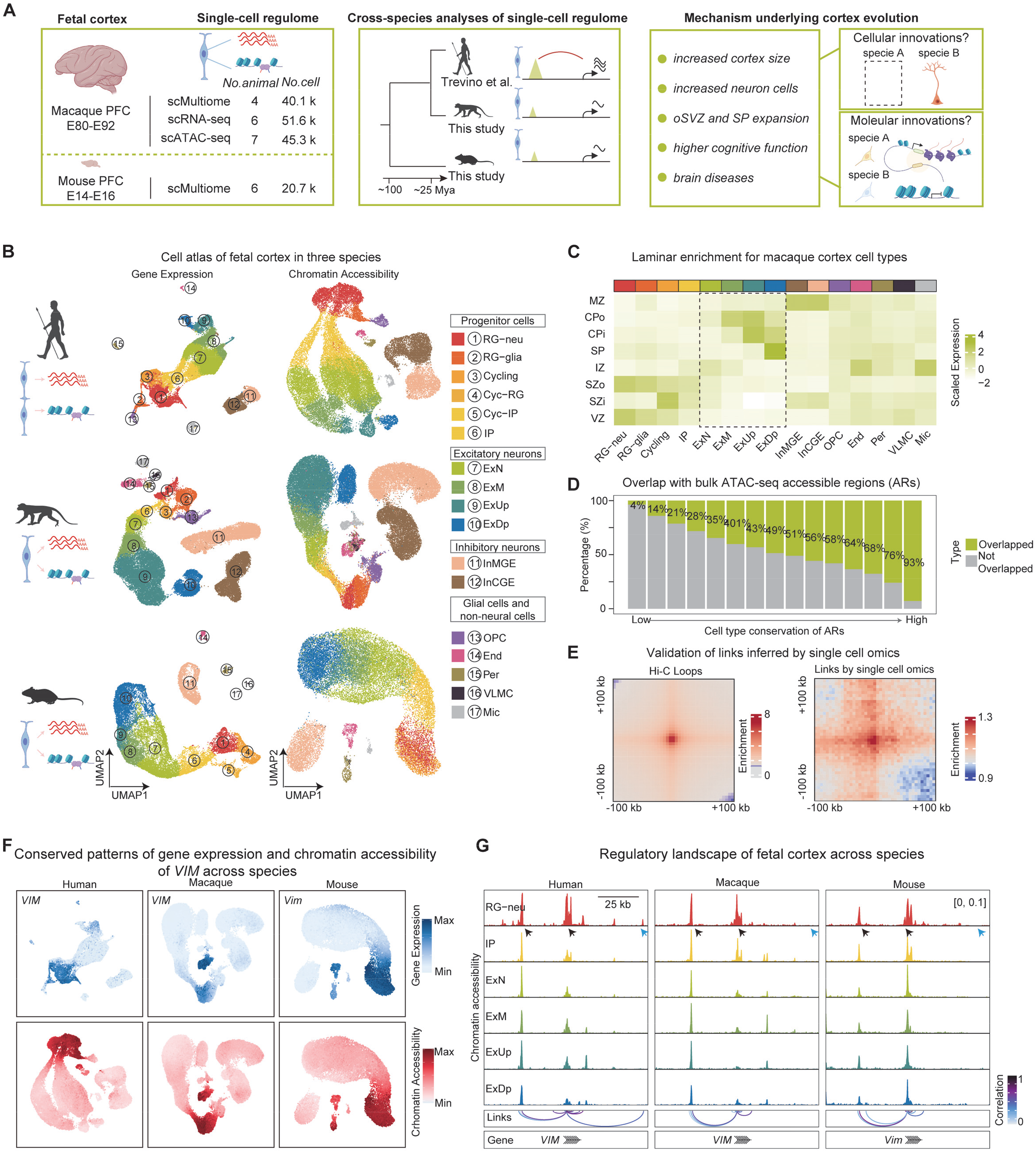
Cell atlas of transcriptome and regulome of the fetal PFC of macaque and mouse during corticogenesis. (A) Schematic view of the study design showing transcriptome and regulome profiling of the fetal PFC of macaque and mouse, and systematic cross-species comparison based on the generated single-cell data. (B) The UMAP plots showing cell clustering patterns (gene expression on the left, and chromatin accessibility on the right) among human (*12*), macaque and mouse. RG-neu, neuron-committed radial glia; RG-glia, glia-committed radial glia; Cyc-RG, cycling radial glia; Cyc-IPC, cycling intermediate progenitor cell; IPC, intermediate progenitor cell; ExN, new-born excitatory neurons; ExM, migrating excitatory neurons; ExUp, upper-layer excitatory neurons; ExDp, deep-layer excitatory neurons; InMGE, medial ganglionic eminence inhibitory neurons; InCGE, caudal ganglionic eminence inhibitory neurons; OPC, oligodendrocytes; End, endothelial cells; Per, pericytes; VLMC, vascular leptomeningeal cells; Mic, microglial cells. (C) The expression patterns of the lamina-specific genes in different cell types of the macaque PFC. marginal zone, MZ; outer cortical plate, CPo; inner cortical plate, CPi; subplate, SP, intermediate zone, IZ, outer subventricular zone, SZo; inner subventricular zone, SZi and ventricular zone, VZ. (D) The overlapped percentages of the chromatin accessible regions (ARs) between scATAC-seq and bulk ATAC-seq. There is a clear correlation between the overlapped percentage and the cell-type conservation of ARs. (E) Consistent enrichment of CRE interactions inferred by the Hi-C data (Hi-C loops) (left) and the scMultiome data (inferred links between expression and ATAC peaks) (right). (F) Conserved patterns of gene expression (snRNA-seq) and chromatin accessibility (snATAC-seq) of the canonical progenitor marker *VIM* among human, macaque and mouse. (G) The overall conserved regulatory profiles around the *VIM* gene region in different cell types among the three species. The conserved peaks are indicated by the black arrows, and the species-specific peak is indicated by the blue arrow.

For evolutionary analyses, as outgroup, we generated the snMultiome data of the mouse PFC at corresponding developmental stages (E14-E16 with 6 biological replicates). The published scRNA-seq and scATAC-seq data of the developmental-stage-matched human PFC (gestation week GW16-GW20) (*12*) were used for cross-species analyses (Fig. 1A). After stringent quality control, we obtained data of 157,799 nuclei, including 40,140 nuclei of macaque snMultiome, 51,623 nuclei of macaque snRNA-seq, 45,327 nuclei of macaque scATAC-seq, and 20,709 nuclei of mouse snMultiome, respectively (Fig. 1A, and fig. S1, A and B). The analyzed developmental stages among the three species fall in the time window of highly active neurogenesis with diverse neural progenitor cells in the developing cortex.

For the macaque PFC data, based on the canonical gene markers, we identified five major cell clusters, including progenitor cells, excitatory neurons, interneurons, glial cells and non-neural cells (Fig. 1B and fig. S1, C to E). The progenitor cells were further classified into four sub-types, including *VIM^+^PAX6^+^EGFR^-^*neuron-committed progenitor cells (RG-neu), *VIM^+^PAX6^-^EGFR^+^*glial-committed progenitor cells (RG-glia), cycling cells and intermediate progenitor cells (IPC) (fig. S1E). Four sub-types of excitatory neurons were identified according to their maturity and laminar distribution, including newborn excitatory neurons (ExN), migrating excitatory neurons (ExM), upper-layer excitatory neurons (ExUp) and deep-layer excitatory neurons (ExDp) (fig. S1E). The laminar distribution of excitatory neurons was further confirmed by the LMD bulk RNA-seq profiles (Fig. 1C and fig. S2A). For example, there are characteristic enrichment of the CP-specific and SP-specific genes in ExUp and ExDp, respectively (Fig. 1C and fig. S2A). Of note, cell annotations are evenly distributed among the macaque biological replicates and highly consistent across data modalities (fig. S1, D and F).

To capture the epigenomic landscape during corticogenesis, based on the macaque scATAC-seq profiles, we detected 67,485-150,000 accessible regions (ARs) per cell type (fig. S2B). Among these ARs, less than 20% are proximal to promoters (within the region of 2,000 bp upstream and 100 bp downstream to the transcription start site) (fig. S2C), suggesting that most ARs may function as distal regulatory elements. Consistent with the known cell-type specificity of distal regulatory elements, about 80% of these ARs were not detected by bulk ATAC-seq and the ARs detected by scATAC-seq showed much higher cell-type specificity compared to ARs shared by the two datasets (Fig. 1D and fig. S2D). Using the correlation method (*15*), we linked the ARs to the putative target genes with matched gene expression and chromatin accessibility profiles. Overall, we identified 49,450 peak-to-gene links involving 40,931 ARs in the macaque cortex (fig. S2E). The enrichment pattern based on the bulk Hi-C profiles validated the accuracy of the inferred peak-to-gene links (Fig. 1E). Combining the single-cell transcriptome and epigenome, we were able to identify cell-type-specific regulations which can be used to define cell identity (fig. S2F). Hence, these findings highlight the utility of single-cell regulome to dissect cell-type-specific regulatory mechanism.

Taken together, the single-cell transcriptome and regulome atlas of the macaque and mouse fetal cortex (Fig. 1, B, F and G, and fig. S3), combined with the previously published human fetal cortex data, allow us to explore cellular innovations and the underlying regulatory mechanisms during primate corticogenesis with a cell-type resolution.

### Evolutionary divergence of progenitor cells in cellular composition

Divergence in cellular composition and properties both contribute to evolutionary innovations (*5*). We first examined the evolutionary changes in cellular composition of neural progenitors during corticogenesis. To evaluate the conservation of cellular architecture in the fetal PFC, we aligned cell types across species based on the gene co-expression pattern (fig. S4). These findings show that the homologous cell types share similar expression patterns and the observed homology is robust to methods and parameters (fig. S4, E and F). Overall, the fetal cortex cell types are highly conserved across species, consistent with the previously reported general conservation of adult cortex cell types (*8*). However, for neural progenitor cells, there are two subclasses showing evolutionary divergence. They are only present in the human and macaque PFC, but absent in the mouse PFC (fig. S4, B, C and F), suggesting a primate specialization in progenitor cell architecture. In light of this result, we conducted a local clustering analysis and re-annotated the neural progenitors of the three species (Fig. 2A). Notably, the proportions of RG-glia and truncated radial glial (tRG) significantly increased in the PFC of human and macaque compared to mouse from early to later gestation stages (Fig. 2B). The expression and chromatin accessibility of the canonical markers, *EGFR* and *CRYAB* for RG-glia and tRG, respectively, confirmed the primate-specific expansion of these two progenitor subclasses (Fig. 2, C and D, and fig. S5, A and B). Moreover, we validated the presence of these two progenitor subclasses using additional published human (*13*) and our macaque PFC datasets with extended developmental stages and data modalities (fig. S5, C and D). Further examination of the mouse cortex dataset (*16*) spanning from E10.5 to E18.5 only detected a very small cluster of RG-glia emerges at E18.5, while the tRG population was extremely rare in all gestation stages (fig. S5E).

**Fig. 2.**
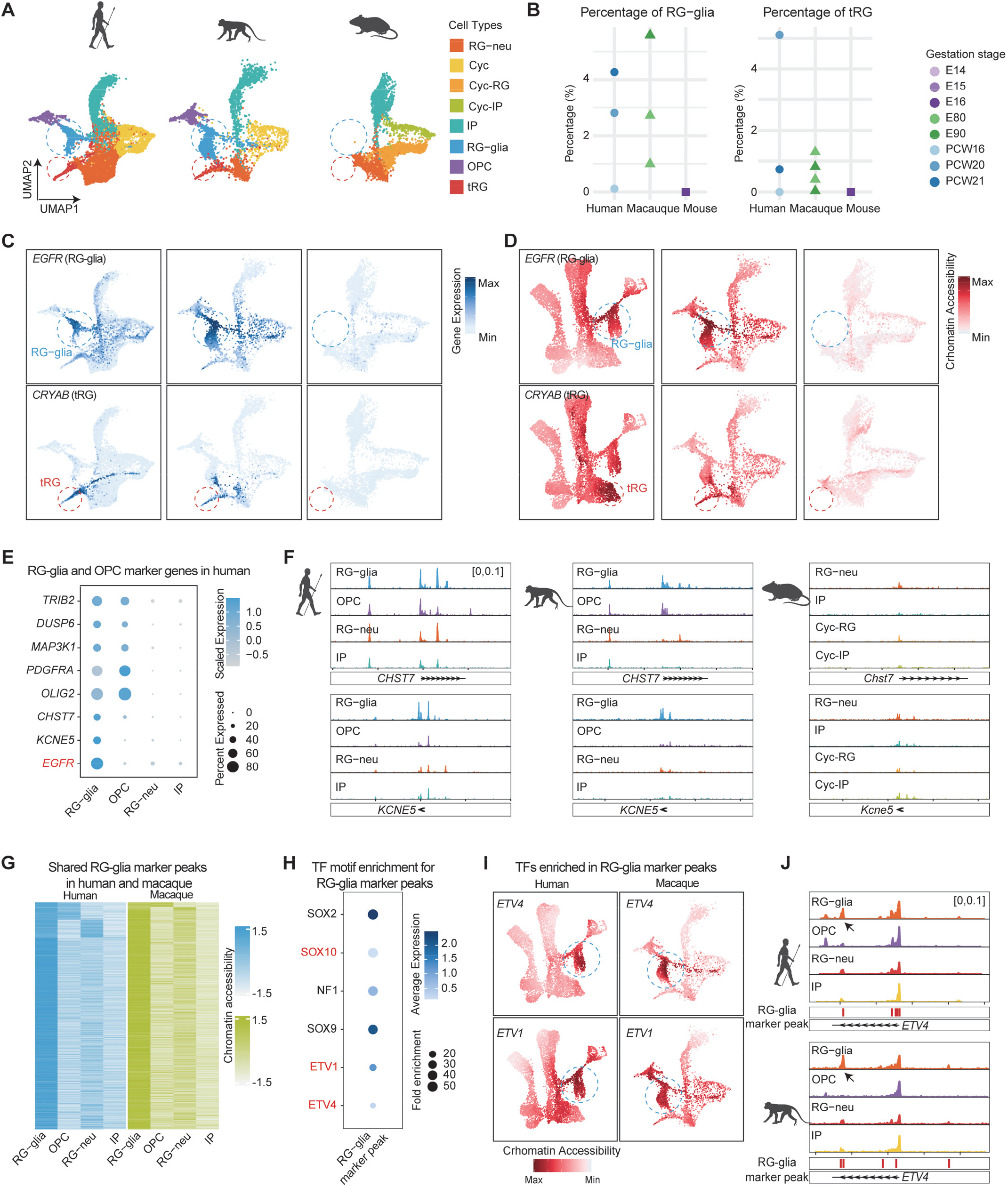
Primate-specific expansion of the RG-glia and tRG cell populations. (A) The UMAP visualization of progenitors among human, macaque and mouse. tRG: truncated radial glia. (B) Comparison of the percentages of RG-glia and tRG relative to the total numbers of the detected cells at different gestation stages. (C) Expression patterns of the canonical marker genes in RG-glia (*EGFR*) and tRG (*CRYAB*). The circled areas indicate the RG-glia (blue) and tRG (red) cell clusters. (D) The distribution of the inferred gene scores of the canonical marker genes in RG-glia (*EGFR*) and tRG (*CRYAB*). The circled areas indicate the RG-glia (blue) and tRG (red) cell clusters. (E) Expression patterns of the identified marker genes in the human RG-glia and OPC. (F) Chromatin accessibility profiles around the two newly identified RG-glia marker genes among the three species. (G) The 1,413 identified RG-glia-specific accessible regions shared between human and macaque. (H) Expression and binding motif enrichment of transcription factors in the RG-glia marker peaks. (I) Gene score distribution for the enriched TFs (*ETV1* and *ETV4*) in the RG-glia marker peaks. The circled areas indicated the RG-glia cell clusters. (J) Chromatin accessibility profiles around *ETV4* in the progenitor cells of human and macaque. The arrows indicated the RG-glia-specific peaks shared between human and macaque.

To gain a deeper characterization of these two progenitor subclasses, we identified their gene markers using our multi-omics datasets. For RG-glia, in addition to known the maker *EGFR*, we identified two new markers, *CHST7* and *KCNE5* (Fig. 2E and fig. S6A). *CHST7* belongs to the galactose/N-acetylgalactosamine/N-acetylglucosamine 6-O-sulfotransferase (GST) family which modulates cell proliferation and cell death signaling pathway (*17*). *KCNE5* encodes a membrane protein of the potassium channel which is implicated in cellular proliferation (*18*). The scATAC profiles of these two genes also displayed signature peaks in RG-glia (Fig. 2F). In addition to these RG-glia marker genes, compared to RG-neu, IP and OPC, we identified 1,413 RG-glia-specific accessible regions shared between human and macaque (Fig. 2G). These RG-glia-specific ARs are enriched in binding motifs for critical transcription factors functioning in neural stem cells and oligodendrocytes, such as SOX2 and SOX10 (Fig. 2H and fig. S6B). Notably, *ETV1* (ETS Variant Transcription Factor 1) and *ETV4* show specific accessibility during oligogenesis (Fig. 2I and J), consistent with a recent study showing *ETV4* as a glial progenitor cell marker (*19*).

For tRG, based on cell type and laminar expression specificity, we detected 13 marker genes including three previously reported markers *CRYAB*, *ANXA1* and *CXCL12* (fig. S6, C to E). All of them showed expression enrichment in VZ of human and macaque based on the LMD RNA-seq data (fig. S6E), consistent with the laminar distribution of tRG in VZ (*20*). The VZ preferential expression of *CRYAB* and *ANXA1* was further confirmed in the human PFC (GW18) using immunofluorescent staining (fig. S6, F and G). Additionally, we detected co-expression of *CRYAB* with three canonical genes (*EOMES*, *MKI67* and *FOXJ1*) in the tRG cells (fig. S6, H and I). These results imply the potential multi-track cell fates of tRG during corticogenesis, and they may give rise to proliferation cells, IPC or endothelial cells. Further examination is required to figure out the exact cell fate of tRG and whether there are differences between human and macaque.

In summary, we discovered primate-specific expansion of RG-glia and tRG cell populations. These results reveal the increased progenitor diversity of primate corticogenesis, which may drive the primate specializations in cortex organization and function.

### Evolutionary divergence of progenitor cells in cellular properties

Besides the increased diversity of cortical progenitors, the primate cortex has specializations in progenitor cellular properties, such as the enhanced self-renewing capacities and the extended duration of neurogenesis (*5*). To systematically dissect the molecular mechanism underlying these evolutionary changes in cellular properties, we searched for human-specific and primate-specific transcriptional programs by analyzing the homologous neuron progenitor cells, RG-neu, which are present in all the studied species.

We used a strict two-step cross-validation method for the cross-species comparison to reduce the noise of single-cell technologies (fig. S7). We first included two additional single-cell transcriptome profiles of the human and mouse cortex to obtain consensus progenitor markers across datasets (*13, 16*) (fig. S7, B and C). Then we compared the chromatin accessibility profiles of the RG-neu progenitor markers across species to detect the inter-specific regulatory divergence (fig. S7D). In total, there are 53 RG-neu markers shared among species, 60 RG-neu markers with primate-specific expression and 74 RG-neu markers showing human-specific expression (Fig. 3A and fig. S7, A and E). Interestingly, only 24% of the human RG-neu marker genes are shared with macaque and mouse, suggesting an evolutionary turnover of molecular signatures defining the homologous cell type, consistent with the reported highly divergent transcriptome of the adult cortex across species (*7, 8*). As expected, the regulatory activities around these markers are consistent with the transcriptional divergence across species (Fig. 3B). For example, the conserved marker *PTN* has similar profiles of chromatin accessibility in RG-neu of all three species. The primate-specific marker *FAM107A* only shows similar profiles in RG-neu in human and macaque, while the human-specific marker *GPX3* displays a human-specific profile (Fig. 3B).

**Fig. 3.**
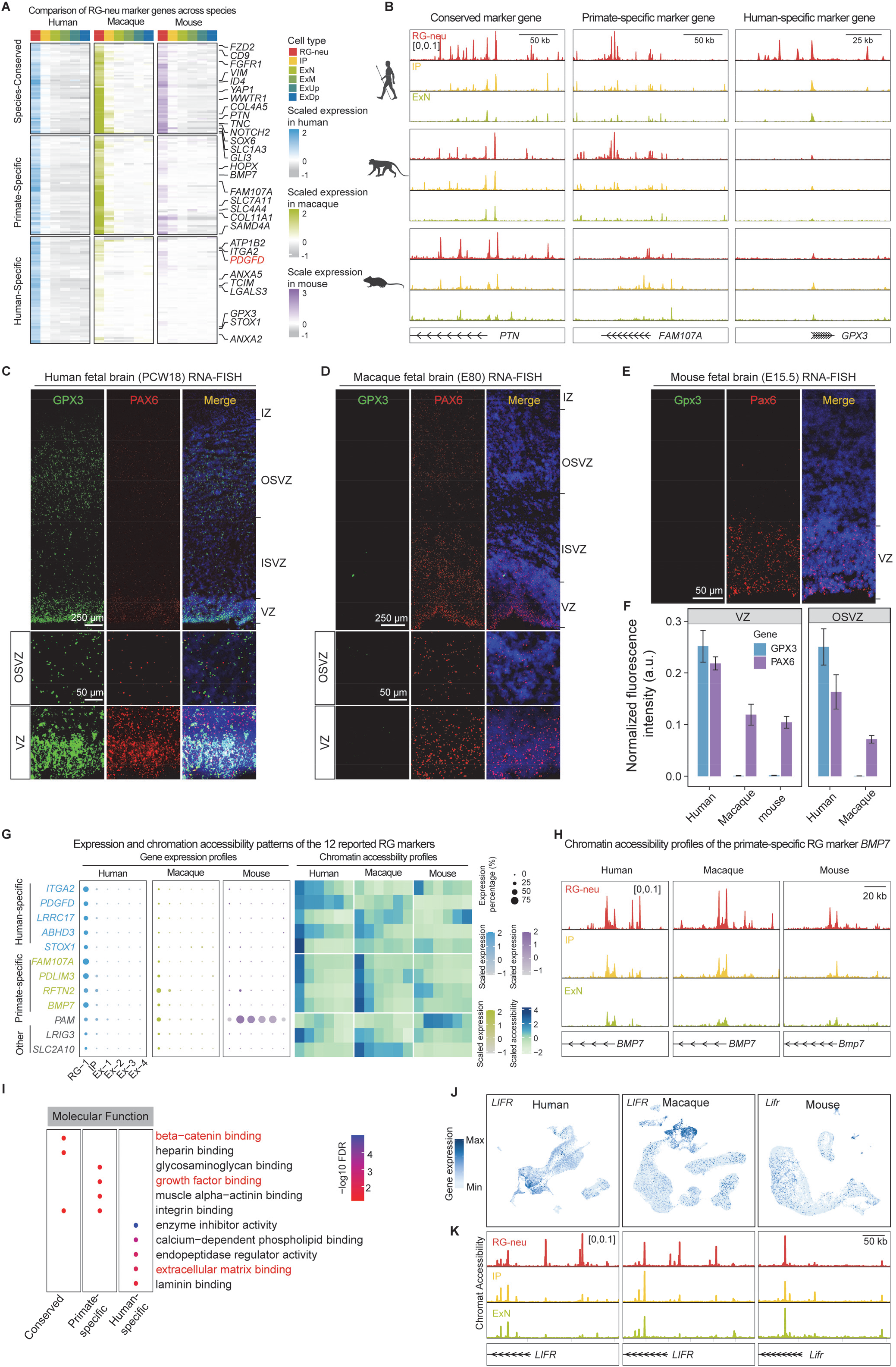
Property innovations of progenitor cells during primate cortex evolution. (A) Identification of conserved, primate-specific and human-specific RG-neu markers. (B) The three gene examples showing conserved, primate-specific and human-specific profiles of chromatin accessibility of the RG-neu markers. (**C-E**) RNA FISH of the human-specific RG-neu marker *GPX3* in the fetal PFC of human (C), macaque (D) and mosue (E). The insets show the human-specific expression of *GPX3* in VZ and OSVZ. (F) Quantification of the normalized fluorescent intensity in VZ and OSVZ for *PAX6* and *GPX3* among human, macaque and mouse. The data is presented in barplot with mean ± SE. The number of the sampled tissue section regions is 33. (G) The expression and chromatin accessibility patterns of the 12 previsouly reported genes with human preferrential expression (human vs mouse) in the RG cells. The data was from Lui et al (*22*). By including the macaque data, 5 human-specific and 4 primate-specific RG-neu markers were identified. (H) The cross-species comparison of the regulatory profiles around the primate-specific marker, *BMP7*. (I) Functional enrichment for the conserved and the species-specific RG-neu markers. The highlighted functional terms in red reflect the potential key mechnisms underlying the conserved and diverged transcriptional wirings in RG-neu. (**J** and **K**) Comparision of the expression profiles (J) and the regulatory profiles (K) of the primate-specific RG-neur marker *LIFR* among human, macaque and mouse.

To confirm the accuracy of the human-specific transcription divergence, we performed single-molecular RNA fluorescence in situ hybridization (smFISH) for the human-specific marker *GPX3*, a gene involved in redox regulation of intracellular signaling (Fig. 3, A and B) (*21*). The results showed a significant high expression of *GPX3* in VZ and OSVZ of the human fetal PFC, and virtually undetectable in the macaque and mouse fetal PFC (Fig. 3, C to F). We compared our species-specific marker sets with 12 previously reported genes showing preferential expression in the human RG compared to the mouse RG (*22*). By including our macaque data, we confirmed that only 5 of them are human-specific (*ITGA2*, *PDGFD*, *LRRC17*, *ABHD3* and *STOX1*), and the others are either primate-specific (4 markers) or not species-specific (3 markers) (Fig. 3G, and fig. 8, A and B). Hence, the macaque data is useful to further narrow down the list of the human-specific marker in RG-neu. Among these genes, *PDGFD* is specifically expressed in the human RG, and is required for human neocortical development (*22*). *BMP7* is identified as a primate-specific RG-neu marker, and it is a secreted growth factor involved in development of bone and kidney (*23*) (Fig. 3H and fig. S8A). Furthermore, using the bulk RNA-seq data (cortical plate and germinal zone of the fetal PFC) (*14, 24*), we validated the up-regulation of the 5 human-specific markers in the human RG-neu compared to the macaque RG-neu (fig. S8 C). These findings demonstrate the accuracy for the identification of species-specific progenitor markers and indicate the utility of non-human primate to dissect human-specific molecular features.

To gain insights into the biological processes associated with primate transcriptional rewiring, we performed gene function enrichment analysis for these marker sets (Fig. 3I, fig. 8D and table S1). The species-conserved markers are enriched in genes associated with Wnt signaling, neuroblast proliferation and focal adhesion regulation (Fig. 3I and fig. 8D). For example, many canonical progenitor markers such as *PTN* and *SOX6* that regulates neurogenesis, as well as the elements for the NOTCH, WNT and YAP signaling pathways (*NOTCH2*, *FZD2*, *ID4*, *WWTR1* and *YAP1*) are shared in RG-neu across species (Fig. 3A), suggesting a general conservation of the core cellular functions of RG-neu. By contrast, genes associated with growth factor binding are significantly enriched in the primate-specific markers (Fig. 3I and fig. 8D). For example, leukemia inhibitory factor receptor (*LIFR*), which was reported to promote self-renewal of neural stem cells (*25*), showed highly specific transcriptional and regulatory signals in the human and macaque RG-neu compared to the mouse RG-neu (Fig. 3, J and K). The growth-factor-associated pathway also shows primate and human specialization during corticogenesis, including the conserved marker *PTN*, the primate-specific marker *BMP7* and the human-specific marker *PDGFD* (Fig. 3A and fig. S8, E and F).

RG cells can generate neurons either directly or indirectly via generation of IPC. We next examined the primate specialization in IPC. As expected, the canonical markers of IPC including *EOMES* and *SMOC1* are shared across species (fig. S9, A and B). Notably, *PPP1R17* displays a primate-specific up-regulation (fig. S9, A to C), in line with the previous report that *PPP1R17* is highly expressed in the primate SVZ but not in ferret or mouse (*26*). Interestingly, there are two HARs (human-accelerated regions) located in the upstream of *PPP1R17*, and the scMultiome profiles suggest their potential involvement in gene regulation (fig. S9D). *PENK* displays a human-specific up-regulation, reflected by both gene expression and chromatin accessibility profiles in IPC (fig. S9, A to C). These findings suggest that gene expression divergence may underlie the diverged proliferation potential in IPC across species.

Together, the cross-species comparison of molecular profiles of neuron progenitors reveals the overall conservation of the core biological processes, and at the same time, there is an extensive rewiring in growth factor and growth factor binding pathways during primate cortex evolution.

### The human-specific ECM-associated gene *ITGA2* promotes progenitor proliferation

In addition to the primate specialization in growth factor pathway, we found that extracellular matrix (ECM) associated genes were significantly enriched in both species-conserved and species-specific RG-neu markers (Fig. 3I and fig. S8D). For example, TNC (Tenascin C), comprised of glycoprotein and highly expressed in the ECM of the central nervous system (*27*), and the collagen family COLL4A5, the most abundant ECM protein, are shared among species (Fig. 4, A and B). *BMP7* and *ITGB5* (encoding a beta subunit of integrin) are specifically expressed in the primate RG-neu (Fig. 4, A and B). Importantly, there are several ECM-related genes with human-specific regulation in RG-neu, including *LGALS3*, *ANXA2* and *ITGA2* (Fig. 4, A and B, and fig. S10A). *LGALS3*, a member of the galectin family, is localized in ECM, and it regulates cell adhesion and progenitor cell proliferation (*28*). *ANXA2* is implicated in modulating cell proliferation and adhesion in tumor (*29*). *ITGA2* (integrin subunit alpha2), which encodes a receptor for collagens and mediates the adhesion of cell to ECM (*30*), marks a niche of epithelial cells in the human placenta (*31*). These results suggest the critical role of ECM pathway in neural progenitor cell function, consistent with previous studies (*32*). At the same time, the ECM-associated pathway has also undergone RG-neu specialization during primate evolution.

**Fig. 4.**
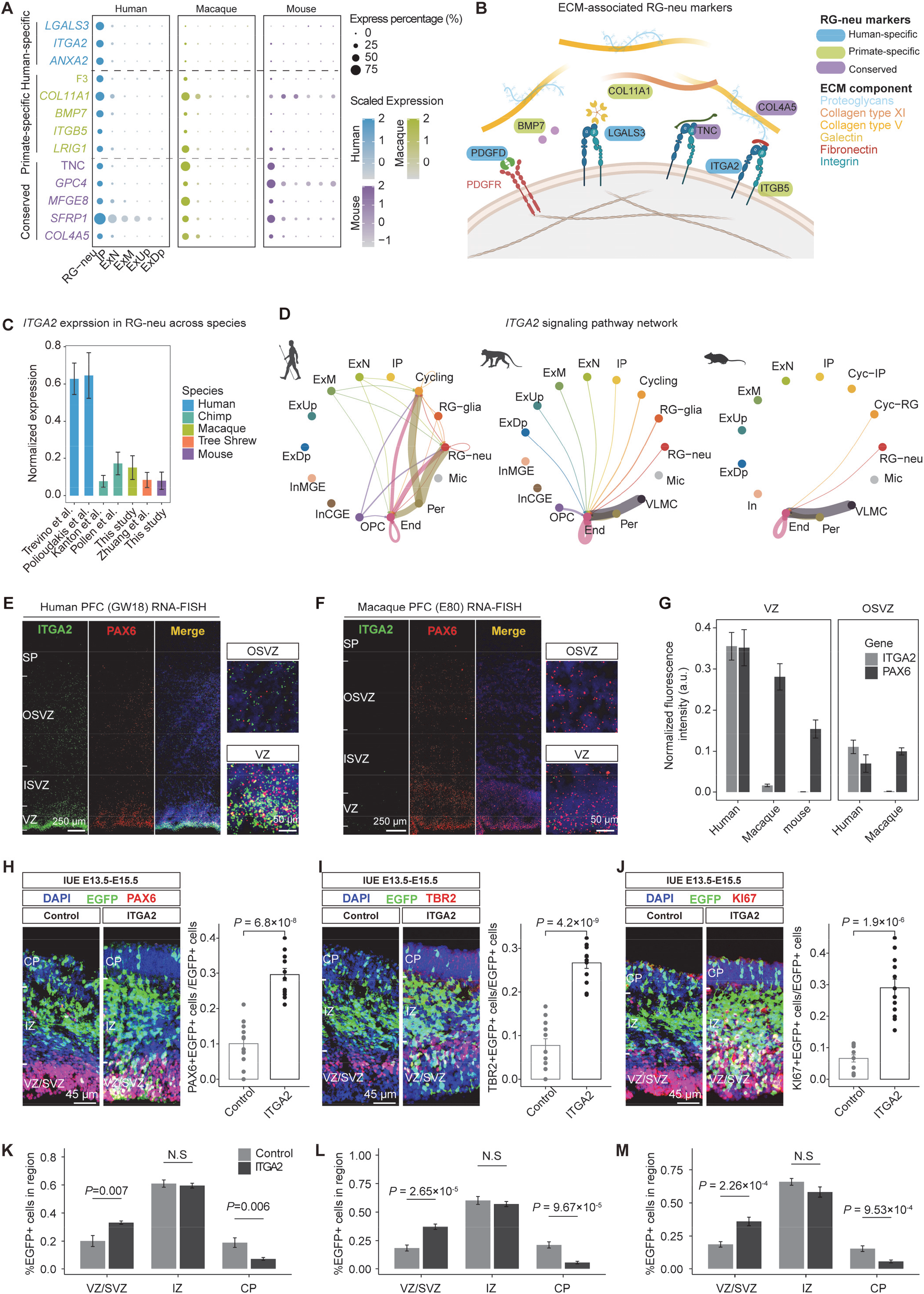
The human-specific changes in the extracellular matrix (ECM) pathway affect neural progenitor proliferation. (A) Expression patterns of the representative conserved and species-specific ECM-associated genes in human, macaque and mouse. (B) Schematic view of the overall structure of ECM and the involved genes. (C) The human-specific high expression of *ITGA2*, compared to chimpanzee (*33, 34*), macaque, tree shrew (*35*) and mouse. (D) The *ITGA2*-associated signaling pathway network indicates the cell-cell communication patterns in the fetal brain. The linewidth indicates the communicating probability. There are clear differences in communication patterns and intensities among the three species. (**E-F**) The RNA FISH visualization of *ITGA2* in the fetal cortex of human (E) and macaque (F). There are abundant *ITGA2*+*PAX6*+ progenitor cells in the human VZ and OSVZ, but rare in macaque. (**G**) Quantifications of the normalized fluorescence intensities of *ITGA2* and *PAX6* in the fetal cortex VZ and OSVZ of human, macaque and mouse. (**H-J**) *In utero* electroporation of *ITGA2*+*EGFP* or *EGFP* only (negative control) in the mouse cortex at E13.5. The fetal cortex was analyzed at E15.5. Co-immunofluorescence-staining of *ITGA2* with three progenitor marker genes, including *PAX6* for neural progenitors (H), *TBR2* for IPCs (I), and *KI67* for cycling cells (J). Compared to the controls, there are significant increases of progenitor cells in the mouse fetal cortex subject to *ITGA2*+*EGFP* electroporation. (**K-M**) Comparisons of the ratios of the *EGFP+* cells in different laminae between the *ITGA2-* electroporation groups and the control groups, correspondent to the samples in (H-J), respectively. The overexpressed *ITGA2* results in more progenitor cells in VZ and SVZ, but not in IZ and CP.

The expression pattern of *ITGA2* caught our attention. It is highly expressed in the human RG-neu, but with much lower expression in chimpanzee, macaque and tree shrew (*Tupaia glis*) (Fig. 4C). Considering *ITGA2* mediates the interaction between cells and ECM (*30*), we examined the cell-cell communications patterns mediated by *ITGA2* among the three species (fig. S10, B and C). There are strong communications involving *ITGA2*, laminin and collagens between the human RG-neu and endothelial/pericytes, but they are much weaker in macaque and mouse (Fig. 4D and fig. S10C). This is in contrast to the similar cell communication patterns across species for the species-conserved RG-neu markers such as *PTN* and *FGFR1* (fig. S10B). Consistently, this human-specific communication pattern was also observed for the other two human-specific RG-neu markers *LGALS3* and *PDGFD* (fig. S11, A and B). These findings indicate that the human specialization in cell communications between progenitor cells and endothelial/pericytes may help maintain the human-specific neural progenitor niche.

We examined the expression pattern of *ITGA2* in the fetal PFC. The smFISH data showed that *ITGA2* is mainly expressed in VZ of the human PFC, with much lower expression in the macaque PFC, and virtually undetectable in the mouse PFC (Fig. 4, E to G, and fig. S11C). We further conducted an *in vivo* electroporation experiment by introducing *ITGA2* into the lateral ventricle of the mouse neocortex at E13.5, and analyzed the fetal cortex at E15.5 (fig. S11D). The results indicated that overexpression of *ITGA2* in the mouse fetal brain led to a significantly increased proportions of progenitors, including RG, IPC and cycling cells (Fig. 4, H to J). Moreover, the overexpressed *ITGA2* seems to help maintain progenitor state of the progenitor cells in VZ and SVZ, but not in IZ and CP (Fig. 4, K to M). Allowing a longer development of the electroporated mouse embryos, at E18.5, we saw a significant increase of upper-layer neurons (fig. S11, E to G). These findings suggest that *ITGA2* can promote neural progenitor proliferation at early embryonic stage, resulting in an increased proportion of upper-layer neurons at later stage.

Previous studies demonstrated that the addition of brain ECM components promotes maturation of human brain organoids (*36*). Consistently, we found that the human-specific ECM-associated genes including *ITGA2*, *LGASL3* and *ANAX2* are significantly up-regulated in the human primary cortex compared to the human brain organoid (fig. S11H). Moreover, the evaluation of the ECM module score also shows a global upregulation in the human primary cortex compared to the organoid (fig. S11I). Consequently, these results suggest that for brain organoid culture, the addition of the human-specific ECM-associated gene products might be beneficial to a better development of brain organoid *in vitro*.

Together, we identified a set of ECM-associated genes showing human-specific specializations in RG-neu, which may contribute to the enhanced proliferation ability of neural progenitors in the developing human brain. Also, this observation provides a hint to optimize the culture conditions of human brain organoid.

### Regulatory mechanism underlying the molecular specializations of the primate progenitors

The detected transcriptional changes in the primate progenitors are presumably attributed to the divergence in activity and interactions of CREs (Fig. 5A). To dissect the regulatory mechanism underlying the molecular changes of the primate progenitors, we sought to compare the chromatin accessibility at single-cell resolution across species per cortex cell type.

**Fig. 5.**
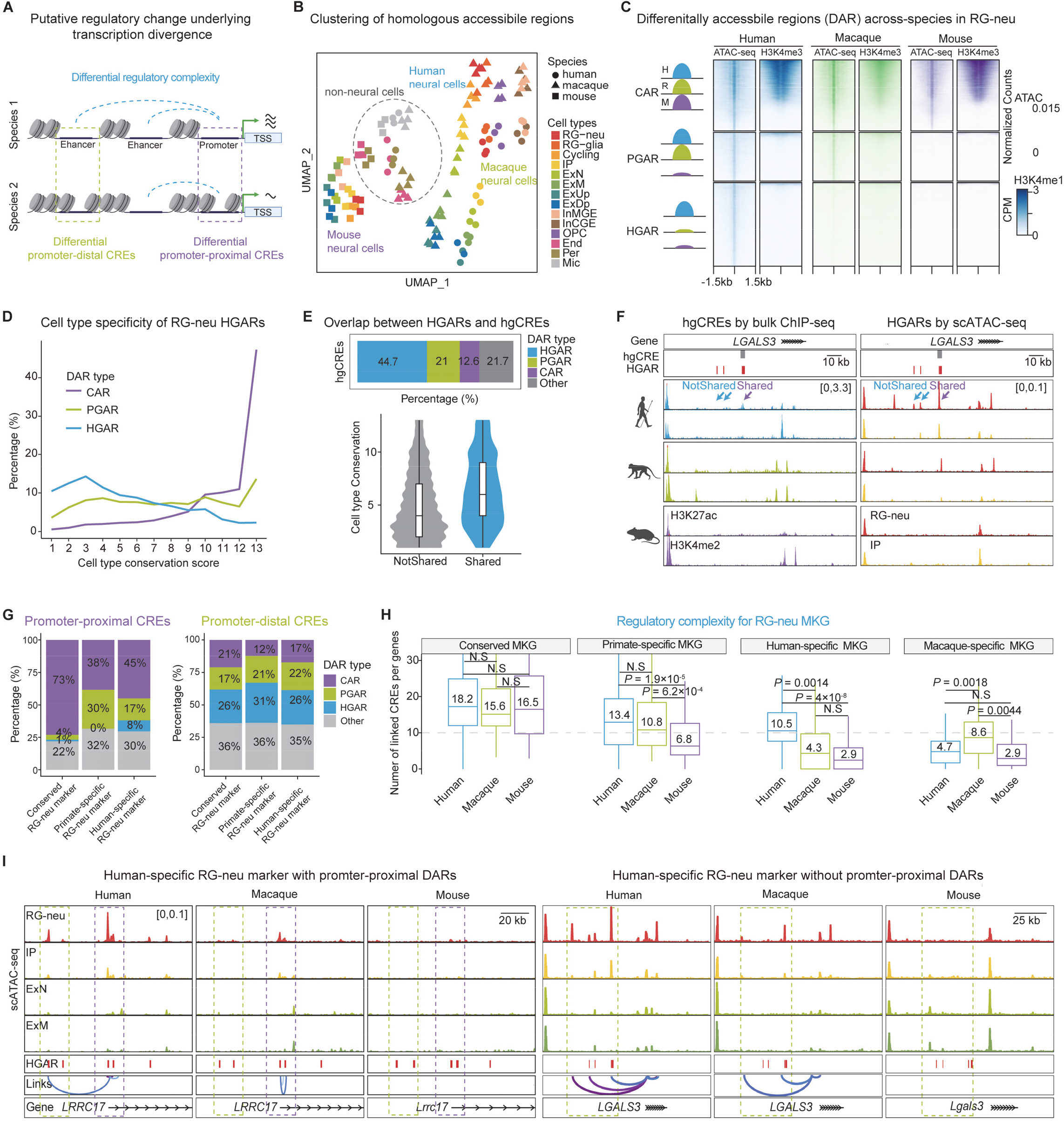
Cell-type-resolved regulatory innovations in the human cortex. (A) Schematic illustration showing changes in gene regulation during evolution. (B) The UMAP plot of the chromatin accessibility profiles for the homologous ARs per cell type among human, macaque and mouse. (C) Heat map showing differentially accessible regions (DARs) across species in RG-neu, including the primate-gained ARs (PGARs) and the human-gained ARs (HGARs), with the conserved ARs (CARs) as reference. (D) Distribution of cell-type conservation scores for CARs, PGARs and HGARs in RG-neu. Conservation score is calculated as the number of human cell types containing ARs per HGAR. (E) Top: Proportions of the previously reported human-gained CREs (hgCREs) (*38*) overlapped with CARs, PGARs and HGARs, respectively. Bottom: comparison of cell type conservation of HGARs that are overlapped or not overlapped with hgCREs. (F) The human-gained ARs located in the *LGALS3* (a human-specific RG-neu marker) gene region, including one shared and two not-shared CREs between HGARs and hgCREs. (G) Percentages of the promoter-proximal and promoter-distal CREs that are overlapped with CARs, PGARs and HGARs among the conserved and species-specific RG-neu markers. (H) Box plots showing the numbers of the linked CREs to the conserved and species-specific RG-neu marker genes. (I) The interspecific divergences in regulatory profiles around the gene regions of *LRRC17* and *LGALS3*.

Chromatin accessibility profiles based on the homologous regions are clustered by species and cell types, and as expected, the neural cells of human and macaque are closely related, and separated from the mouse neural cells (Fig. 5B). To identify the genomic regions showing human-specific gains of chromatin accessibility, we quantified the chromatin accessibility changes at the homologous regions across species using single-cell and pseudobulk sampling strategies (fig. S12, A to D, and table S2 to S4) (*15, 37*). Overall, based on the cross-species comparison, we identified 4,838-21,785 human-gained accessible regions (HGARs), 1,994-11,748 primate-gained accessible regions (PGARs) and 8,335-18,631 conserved accessible regions (CARs) per cell type (Fig. 5C and fig. S12, A and B). We showed that the identified HGARs were not method-biased, and the two HGAR sets by two different methods are highly consistent (fig. S12C).

To explore the regulatory function of HGARs, we first examined their genomic distribution. It turns out that in RG-neu and IPC, there are only 13% of HGARs being proximal to promoters, contrasting 67% in CARs, indicating that the great majority of the human-specific CRE rewiring are probably distal (Fig. 5C and fig. S12E). Consistently, HGARs show a higher cell-type-specificity compared to CARs (Fig. 5D and fig. S12F). We reckoned that most HGARs would not be captured using the regulatory profiles of bulk samples. Indeed, when comparing the identified HGARs with the previously reported human-gained CREs (hgCREs) using bulk samples (*38*), we saw a much higher cell-type specificity in those HGARs not shared with hgCREs (Fig. 5E and fig. S12G). For example, the gene region of *LGALS3* (one of the three identified ECM-associated human-specific RG-neu markers) contains three HGARs, and only one of them is detected in the bulk sample inferred hgCREs (Fig. 5F). These results demonstrate the utility of single-cell regulome in uncovering cell-type-specific regulatory innovations in the human cortex.

We also investigated the evolutionary origin of HGARs per cell type. The majority of HGARs (76%-80%) are likely generated through modification or co-option of the ancestral regulatory elements since they are shared among the studied species. Remarkably, there are 20%-26% *de novo* HGARs that are absent in macaque and mouse (fig. S12, H and I). These *de novo* HGARs likely originated from the non-regulatory sequences in the genome, and may have a significant contribution to the human-specific regulatory innovations.

Next, we explored the regulatory changes orchestrating the human-specific progenitor identity genes. Putative distal CREs regulating maker genes were inferred based on the peak-to-gene links (fig. S13). The transcriptional divergence is primarily associated with the changes of distal CRE activities, and only 17% of the distal CREs linked to the human-specific markers are conserved across species, while the ratio of the conserved promoter-proximal CREs is 45% (Fig. 5G and fig. S12J). Moreover, the human-specific RG-neu markers show significantly higher regulatory complexity (reflected by the number of the linked CREs per gene) than in macaque and mouse (Fig. 5H). For example, *LRRC17* and *LGALS3* are both human-specific RG-neu markers with upregulation in the human RG-neu compared to macaque and mouse. The interspecific transcriptional divergence of *LRRC17*is likely driven by both promoter-proximal and promoter-distal CREs, while for *LGALS3*, only the activity changes of the promoter-distal CREs were detected (Fig. 5I).

Together, utilizing the single-cell regulatory profiles, we identified human-gained CREs associated with the regulatory changes underlying molecular innovations of progenitor cells.

### Genetic basis and neuropsychiatric relevance of the human regulatory innovations

To identify genetic basis of the human specialization during corticogenesis and the increased vulnerability to neuropsychiatric disorders (Fig. 6A), we first evaluated the enrichment of the human-specific sequence changes in the HGAR regions. As expected, HGARs are significantly enriched with the human-specific single nucleotide changes (SNCs) compared to CARs and PGARs (Fig. 6B). Given HARs contain multiple human-specific SNCs and are thought to drive human-specific gene regulations, we screened out those HGARs that overlapped with HARs. Similar to the enrichment seen in the human-specific SNCs, we found significant enrichment of HARs in HGARs among the three reported HAR sets identified by different methods (fig. S14A) (*26, 39, 40*). In total, there are 359 HAR-overlapping HGARs (fig. S14, B and C). Moreover, HGARs show higher enrichment in HARs than CARs in the majority of the cortex cell types, including RG-neu, ExM, ExUp and InMGE (Fig. 6C).

**Fig. 6.**
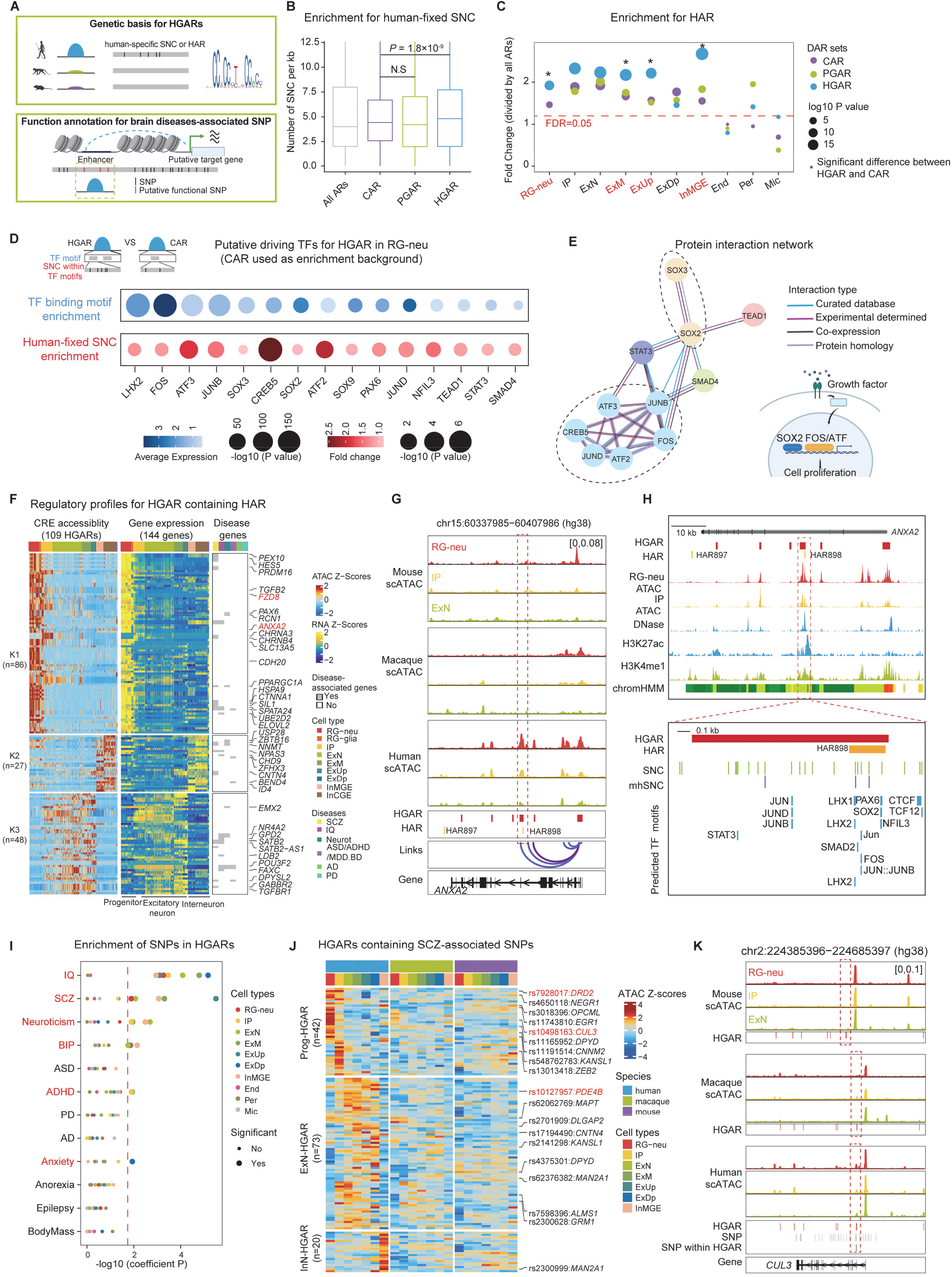
Genetic basis and neuropsychiatric relevance of the human regulatory innovations. (A) Schematic illustration showing the genomic basis and implication in brain diseases for HGARs. (B) The density of the human-fixed single nucleotide changes (SNCs) indicates the enrichment of these SNCs in HGARs compared to CARs in RG-neu. P values are calculated by T test. (C) Enrichment of the human-accelerated regions (HARs) in the conserved and species-specific ARs per cell type. Fold change is calculated by dividing the HAR-overlapping percentage for CARs, PGARs and HGARs by the HAR-overlapping percentage for all homologous ARs. P values are calculated by Fisher’s exact test. The dashed line indicates the significance threshold for Benjamini-Hochberg adjusted P value. Four cell types (RG-neu, ExM, ExUp and InMGE) showing significant differences between HGARs and CARs are highlighted in red. (D) The inferred transcription factors with detectable expression in the human RG-neu show binding motif enrichment in HGARs, as well as enrichment of the human-specific SNCs in these TF motifs. (E) Left: the protein interaction network for the inferred transcription factors from (D) based on STRING database. Right: the schematic illustration showing the possible signaling associated with these transcription factors. (F) Heat maps showing chromatin accessibility of HGARs overlapped with HARs (left), the expression of the predicted target genes (middle) and the related brain diseases (right). (G) The example of HGAR overlapped with HAR around the *ANXA2* gene region. The normalized mulitome scATAC-seq tracks in human, macaque and mouse are shown. The HGAR containing HAR898 is indicated by the red box. (H) Top: The normalized tracks covering mutiome scATAC-seq, DNase-seq, H3K27ac, H3K4me1 and the predicted chromatin states in the human fetal brain. Bottom: A zoom-in view of the HGAR overlapped with HAR898. The human-fixed SNCs and the predicted TF binding motifs are indicated. The data of DNase-seq, H3K27ac and H3K4me1 were from the previous studies (*50, 51*). The modern-human-fixed SNCs (mhSNC) were from the previous study (*52*). (I) Heritability enrichment of the brain diseases-associated GWAS SNPs in HGARs per cell type based on linkage disequilibrium score regression (LDSC). The dashed line indicates the threshold for Benjamini-Hochberg adjusted LDSC coefficient P value. (J) Heat maps showing chromatin accessibility of the HGARs overlapped with the SCZ-associated SNPs in human, macaque and mouse. The SNP IDs and their nearest genes are labeled. (K) The example of the RG-neu HGAR overlapped with the SCZ-related SNPs (indicated by dashed box) around the *CUL3* gene region. The normalized multiome scATAC-seq tracks are shown.

To look for the putative transcription factors involved in the human-specific regulatory changes in RG-neu, we screened out a set of candidates with detectable expression in the human RG-neu, preferential binding motif enrichment and human-specific SNC enrichment in HGARs compared to CARs (Fig. 6D and fig. S14, D and E). These TF candidates include the known TFs critical for neural progenitors such as LHX2, SOX2 and PAX6. Notably, there is a cluster of TFs that shows protein sequence homology including FOS, the JUN family, and the ATF family (Fig. 6E), and they are thought to response to environmental stimuli such as growth factor that can activate cell proliferation (*41, 42*). The binding motifs of these TFs are more accessible in the human RG-neu compared to macaque and mouse (fig. S14F), indicating their potential roles in shaping the human-specific regulatory activities. In addition, *FOS* has a primate-specific expression in VZ and SVZ (fig. S14G), an indication of a previously unappreciated role of this gene in neural progenitors.

We also identified 144 putative target genes among the109 HAR-overlapping HGARs using the peak-to-gene approach (Fig. 6F and table S5). Importantly, more than half of these HARs show progenitor-specific accessibility and their putative target genes are specifically expressed in the human progenitor cells (Fig. 6F). In additional to the previously reported putative HAR target gene *FZD8* (*43*) (Fig. 6F), many other genes critical for cortex development are also predicted to be regulated by HARs such as *PAX6*, *HES5* and *ID4* (Fig. 6F). Notably, *ANXA2*, a gene with a human-specific expression in RG-neu, is overlapped with HAR898 that shows activity gains in the human RG-neu compared to macaque and mouse (Fig. 6, G and H, and fig. S14, H to J). More importantly, HAR898 exhibits an enhancer activity based on the epigenomic profiles of the human fetal brain, and it harbors 5 human-specific SNCs that are predicted to introduce novel transcription factor bindings including FOS, LHX2 and SOX2 (Fig. 6H).

Finally, we asked whether the inferred regulatory innovations for human cortex development would increase our vulnerability to neuropsychiatric disorders. Of note, many of these putative HAR target genes are associated with cognitive traits and early-onset brain diseases, such as IQ, SCZ and autism spectrum disorder (ASD) (Fig. 6F) (*44*). Given these associations, we investigated the links between HGARs and the brain-related traits or diseases. We first performed linkage disequilibrium score regression (LDSC) to evaluate the enrichment of single nucleotide polymorphisms (SNPs) in human populations associated with brain-related traits and diseases per cell type in HGARs (Fig. 6I). The results indicate that HGARs have significant enrichment in multiple traits and early-onset diseases including IQ, SCZ, neuroticism, bipolar disorder (BIP), attention-deficit/hyperactivity disorder (ADHD) and anxiety in at least one cortex cell types (Fig. 6I). Similarly, a recent study demonstrated that the human fetal dorsolateral prefrontal cortex-specific CREs (compared to cerebellar cortex) are enriched in IQ, SCZ, ASD and neuroticism (*44*). These findings suggest that during the emergence of the human-specific CREs, due to linkage, genetic hitchhiking would cause the elevation of the risk SNCs in human populations, leading to an increased vulnerability of neuropsychiatric disorders.

Markedly, IQ- and SCZ-associated SNPs showed the highest enrichment scores (Fig. 6I). A total of 133 HGARs contain SCZ-associated SNPs, and these SNPs are located in many known risk genes for SCZ (Fig. 6J). For example, the D2 dopamine receptor coding gene (*DRD2*) is one of the most important candidate genes for SCZ and also the therapeutic target for neuropsychiatric disorders (*45, 46*). *PDE4B*, which shows translocation in the SCZ patients, regulates cAMP signaling that is implicated in learning, memory and mood (*47*). Another noticeable example is *CUL3*, a high confidence risk gene for autism, schizophrenia and developmental delay (*48, 49*). The gene body of *CUL3* contains many SCZ-related SNPs and one of them shows human-specific accessibility in RG-neu, supporting the speculated genetic hitchhiking effect in HGARs that increases vulnerability of neuropsychiatric disorders (Fig. 6K).

Altogether, the identification of HGARs at cell-type resolution pinpoints candidate casual genetic changes responsible for the human specializations during corticogenesis and the increased risk for neuropsychiatric disorders, an indication of the interplay between brain evolution and brain diseases.

## Discussion

We generated a comprehensive regulatory landscape of the macaque and mouse fetal PFC at single-cell resolution. Through integrative analysis with the published human fetal PFC data (*12, 13*), we demonstrated the human specializations in neural progenitor cellular architecture and properties, and deciphered the underlying regulatory mechanism.

The human cortex expansion relies on divergence in progenitor cell organization and functions that reside in VZ and SVZ (*4, 5*). Although cortex cell types are overall conserved across species during corticogenesis, we found two progenitor subtypes that are significantly expanded in the primate cortex including RG-glia and tRG, highlighting the role of cellular innovation on progenitor diversity during primate cortex evolution. These primate-specific progenitors have been reported to give rise to the glial lineage including oligodendrocyte precursor cells (OPCs) and astrocytes in the human cortex (*19, 53*). This is in line with the recently reported human-specific increase of OPC compared to NHPs and mouse (*11, 54*). Therefore, this primate-specific expansion suggests an extensive overlapping between neurogenesis and gliogenesis during primate cortex development, and indicates the interplay between cortex expansion and white matter expansion during primate cerebral evolution.

Furthermore, we identified primate specializations in neural stem cell niches including the growth factor- and ECM-associated pathways. In particular, we found primate-specific expression of growth factor *BMP7* and LIF receptor and human-specific expression of *PDGFD* in neural progenitor cells. The ECM-associated genes including *ITGA2*, *LGALS3* and *ANXA2* exhibit human-specific gains of CREs that upregulate their expression in the human progenitor cells compared to macaque and mouse. This is in line with the previous report that the ECM pathway is enriched in genes associated with the human-gained CREs (*38*). The ECM is implicated in regulating the proliferative potential of cortical progenitor cells in human and mouse, likely via forming a stem cell niche with local growth factors (*27, 32*). The ECM components also participate in the cellular mechanotransduction and possibly induce cortical folding (*36, 55*). Specifically, a recent study demonstrates that the binding protein (LGALS3BP) of LGALS3 regulates neural progenitor anchoring and migration, and its mutations result in altered gyrification and cortex thickness in patients (*56*). Of note, these ECM-associated genes were down-regulated in human brain organoid compared to primary human cortex. Hence, these findings indicate the human-specific and primate-specific modifications in the neural stem cell niches compared to rodent during corticogenesis, which may account for the enhanced progenitor proliferation in the primate (especially human) cortex. These newly discovered genes with human and primate specializations shed light on the molecular mechanism of cortex evolution, and at the same time may help improve the culture conditions of human brain organoids.

Changes in gene regulation are the key driving force of evolution (*2*). Based on the comprehensive regulatory mapping of the macaque PFC during corticogenesis, we explored the regulatory mechanism underlying the human specializations in progenitor gene expression programs. We found that the expression divergence is mostly driven by the activity changes of distal CREs, which is in line with the reported rapid evolution of enhancers (*38, 57*) and our previous observation of enriched enhancer-enhancer interactions in the human-specific chromatin loops (*14*). We also identified thousands of HGARs per cortex cell types. HGARs are mostly cell-type specific, indicating context-specific functional innovations for the regulatory divergence in the developing human cortex. Moreover, HGARs show significant enrichment in the human-specific genetic changes including HARs. Linking the genetic changes to the human cortex specialization is challenging due to the lack of comprehensive functional interpretations of the fetal cortex of closely related species such as NHPs. With the identification of HGARs per cell type, we provided a candidate list of causal human-specific genetic changes. Further functional validations for these HGARs will aid the dissection of genetic mechanism underlying human cortex evolution. In addition, the discovery of conserved and species-specific accessible regions can be useful in designing the labeling tools targeting certain cortex cell types.

Finally, we also observed enrichment of HGARs in SNPs associated with IQ and early-onset brain disorders including SCZ, neuroticism and bipolar disorder. In particular, several high-risk genes and well-established drug targets are regulated by HGARs, such as *DRD2*, *PDE4B* and *CUL3*. These findings indicate the interplay between cortical development and neuropsychiatric disorder via gene regulation, and this connection may reflect the evolutionary trade-offs of the human neuropsychiatric vulnerabilities. This is in line with the previously reported convergence in early onset neuropsychiatric disorders for genes associated with the human-specific regulatory changes (*38*), and our results pinpointed the fetal brain cell types susceptible to these brain diseases. Taken together, the generated comprehensive regulatory landscape at single-cell resolution for the macaque and mouse fetal cortex, combined with the evolutionary dissection of the human cortex, provides a rich resource for future studies on primate brain development, evolution and diseases.

## Acknowledgments

We thank Yan Guo for their technical assistance in this study. We thank Changjie Sun from Kunming Institute of Zoology for his help in IUE experiment. Part of the analysis was performed on the High Performance Computing Platform of the Center for Life Science of Peking University.

## Author contributions

Conceptualization: Y.L., X.L., C.L., B.S.

Methodology: Y.L., X.L., Y.S., K.C., H.T., B.Y., J.X., F.Z., X.M., X.L. and X.H.

Investigation: Y.L., X.L., Y.S. and K.C.

Visualization: Y.L., X.L., Y.S. and K.C. Funding acquisition: B.S., C.L.

Project administration: Y.L., X.L., C.L. and B.S. Supervision: X.L., C.L. and B.S.

Writing – original draft: Y.L., X.L. and Y.S.

Writing – review & editing: Y.L., X.L., Y.S, C.L. and B.S.

## Funding

This study was supported by grants from National Natural Science Foundation of China (U2002207, 32000406 and 32170628), National Science and Technology Innovation 2030 Major Program (STI2030-2021ZD0200100), and the Youth Innovation Promotion Association of CAS (to X.L.). Y.L., Y.S. and C.L. were supported by National Natural Science Foundation of China (32288102, 32025006) and National Key Research and Development Program of China (2021YFA1100300).

## Competing interests

The authors declare that they have no competing interests.

## Data availability

The snMulitome data for macaque and mouse PFC, snRNA-seq, scATAC-seq, LMD bulk RNA-seq and bulk ATAC-seq data were deposited at NCBI GEO under the accession number: GSE241429.

## Code availability

Scripts used in this study are available at Github repository: https://github.com/yimingsun12138/BrainEvoDevo. All data are available in the main text or the supplementary materials.

## Methods

### Animals

Fetal brains of rhesus macaque (embryonic day 80-92) were obtained from Kunming Primate Research Center, Kunming Institute of Zoology. The C57BL/6J mice at embryonic day 14-16 were used for single-cell multiome libraries construction and RNA FISH. Pregnant ICR mice (embryonic day 13.5-18.5) were used for in utero electroporation and immunostaining studies. All C57BL/6J and ICR background mice were obtained from Animal Center, Kunming Institute of Zoology. All animal procedures were conducted following the international standards, and were approved in advance by the Institutional Animal Care and Use Committee of Kunming Institute of Zoology, Chinese Academy of Sciences (Approval No: SMKX-SQ-2021-091).

### Human fetal brain samples

The human fetal brain (GW18) was used for RNA FISH and immunostaining studies. The brain tissue collection and research protocols were approved by the Reproductive Study Ethics Committee of Nanjing Maternity and Child Health Care Hospital (ethics committee).

### Nuclei isolation

Transfer fresh frozen tissue sample to a pre-chilled 1.5 mL Eppendorf tube, add 500 μL of chilled Nuclei Lysis Buffer, and then transfer to a Douncer. Dounce the tissue with A and B pestle for complete homogenization, filter with a 40 μm strainer, and add 500 μL of chilled Nuclei Lysis Buffer. Mix gently and incubate on ice for 5 min. Centrifuge the nuclei at 500 x g for 5 min at 4°C. Remove supernatant without disturbing the pellet. Isolate pure nuclei using gradient centrifugation, and quantify the nuclei using Trypan blue staining on a hemocytometer.

### Library constructions of snRNA-seq, snATAC-seq and scMultiome

For single-nucleus multi-omics library construction, the isolated nuclei were resuspended with 500 μL diltuted nuclei buffer (1 x nuclei buffer (10X Genomics, PN-2000207), 1mM DTT, 1U/μL RNase inhibitor, nuclease-free Water). The nuclei were counted using Trypan blue staining on a hemocytometer, and 6,000 nuclei were loaded per lane. All libraries were constructed based on the user guide of Chromium Next GEM Single Cell Multiome ATAC + GEX kit (10X Genomics). Libraries were sequenced on the Illumina NovaSeq 6000 platform.

### Data processing of the macaque LMD bulk RNA-seq

The raw sequencing reads were processed by ‘fastp’ to trim adapters. The Mmul_10 reference genome and the macaque GTF gene annotation file obtained from Ensembl (release-103) were used to construct the STAR indexing (v2.7.9a) (*58*). Paired-end sequencing reads were then mapped to the reference genome. The SAM files were converted to BAM format using ‘samtools’ (v1.16.1) (*59*). Feature counts were calculated using the ‘summarizeOverlaps’ function from the ‘GenomicAlignments’ R package (v1.34.0). Laminae-enriched genes were detected by ‘DESeq2’ (v1.38.3) (*60*) with parameter settings “design = ∼ sample source + laminae, log2FoldChange threshold = 0, FDR threshold = 0.05”.

### Data processing of the macaque bulk ATAC-seq

Low quality reads and adapters were filtered by ‘fastp’ with default parameters. The filtered reads were aligned to the Mmul_10 reference genome using ‘Bowtie2’ (v2.5.0) (*61*). Duplicated reads were removed by ‘picard’ (v2.27.5). Only uniquely mapped alignments (MAPQ > 30) were retained for further analysis. Peaks were detected using ‘MACS2’ (v2.2.7.1) (*62*) with settings “callpeak -q 0.01 –nomodal --extsize 147”.

### Data processing of the macaque snRNA-seq

The custom macaque reference genome was constructed by employing the same genome and genome annotation file mentioned in macaque bulk RNA-seq processing, using the command ‘cellranger mkref’ (v6.0.0, 10x Genomics). The sequencing reads were aligned to this reference genome using the command ‘cellranger count --include-introns’. The resulting gene expression count matrix was then introduced into the ‘Seurat’ R package (v4.3.0) (*63*) for further analysis. Cells with fewer than 200 detected genes were filtered out as low-quality data and cells with either total reads greater than 15000 or detected genes greater than 7000 were filtered out as doublets. Next, we normalized the gene expression count matrix by dividing the total reads detected in each cell, scaling to 10000, and then applying a log transformation. Using the ‘vst’ algorithm built in ‘Seurat’, we identified 3000 most variable genes among cells and then reduced the gene expression count matrix to 50 dimensions through Principal Component Analysis (PCA) based on these genes. The top 31 principal components that contributed the most variation were selected for further analysis. Shared Nearest Neighbor (SNN) graph was constructed using the ‘FindNeighbors’ function. Clusters were identified by applying Louvain algorithm to the SNN graph, using the ‘FindClusters’ function with the resolution parameter set to 1. To further remove doublet cells, we utilized the ‘scrublet’ Python package (v0.2.3) to calculate a doublet score for cells in each sample. Clusters with more than 80% of cells predicted to be doublets were removed. Finally, we employed the ‘RunUMAP’ function with default parameters to generate a two-dimensional representation of the 52924 cells that passed the filter.

### Data processing of the macaque snATAC-seq

The same reference genome and genome annotation file as mentioned above, along with JASPAR2022 vertebrate core PFMs, were used to construct the macaque custom reference genome, using the commond ‘cellranger-atac mkref’ (v2.0.0, 10x Genomics). Subsequently, snATAC-seq reads were mapped to this reference genome, using the command ‘cellranger-atac count’. The fragment files were analyzed using the ‘ArchR’ R package (v1.0.2) (*15*). To ensure high-quality data, we filtered out cells with fewer than 1000 fragments or TSS enrichment scores less than 4. For low-dimensional representation of snATAC-seq data, we employed the iterative Latent Semantic Indexing (LSI) method provided by ‘ArchR’. Specifically, the whole genome was divided into tiles of 5kb length. In the first iteration, the most accessible 20000 genome tiles were used for LSI reduction. Clusters were identified using the Louvain algorithm (integrated in ‘Seurat’) based on the LSI reduction, with the resolution parameter set to 0.6. In the second iteration, the accessibility was summed across cells in each cluster, and the most variable 20000 genome tiles were used for LSI reduction and clustering as mentioned above. In the final iteration, the top 500000 most variable genome tiles, based on the clustering in the second iteration, were selected for the final LSI reduction. To correct for the batch effect caused by the experiment, the LSI reduction was adjusted using the ‘addHarmony’ function integrated in ‘ArchR’. The top 25 singular values were then used for final clustering and UMAP visualization with the resolution parameter in the Louvain algorithm set to 1, and the minimal distance parameter in UMAP set to 0.6.

To achieve accurate and robust cell type annotation, we used both marker gene activity scores and integration with snRNA-seq data to annotate each cell in snATAC-seq data. Gene activity scores represent the accessibility around TSS site for each gene and were calculated using the ‘addGeneScoreMatrix’ function with default parameters. Due to the sparsity of the snATAC-seq data, we employed the ‘MAGIC’ method available in ‘ArchR’ to impute gene activity scores by smoothing signals across nearby cells. Using the gene activity score matrix, we initially assigned a rough category for each cell in the snATAC-seq data, including neuron progenitor cells, excitatory neurons, inhibitory neurons and other cell types. We then performed constrained integration between macaque snRNA-seq and snATAC-seq data, using the ‘addGeneIntegrationMatrix’ function.

After carefully annotation, cells from each cell type were sampled into different pseudo-bulk replicates matching their respective sample source. The ‘addReproduciblePeakSet’ function and ‘MACS2’ software (v2.2.7.1) (*62*) were used for peak calling on the aggregated insertion sites in each pseudo-bulk replicate (q value cutoff was set to 0.05), and only reproducible peaks were retained. To create a consensus set of peaks, an iterative overlap peak merging procedure integrated in ‘ArchR’ was performed to merge the peak sets from all cell types.

### Data processing of the macaque snMultiome

The custom macaque reference was constructed with the same files mentioned above, with the commond ‘cellranger-arc mkref’ (v2.0.0, 10x Genomics). Raw sequencing files were aligned to this custom reference genome and quantified using the command ‘cellranger-arc count’. The transcriptome modality was analyzed using ‘Seurat’, while the chromatin accessibility modality was analyzed using ‘ArchR’ following the same standards described above. We used the same criteria to filter low-quality data and doublets in both gene expression matrix and chromatin accessibility data. Similar clustering and annotation were performed on the transcriptome modality as described in macaque snRNA-seq processing, with only a few parameter changes (top 31 principal components were used for clustering and UMAP visualization, and the Louvain clustering resolution was set to 0.7). Cell type annotation in the chromatin accessibility modality was inherited from the transcriptome modality, followed by UMAP visualization (top 14 singular values were used) and peak calling.

### Data processing of the mouse snMultiome

The mouse custom reference genome was constructed by employing the GRCm38 genome assembly and the mouse GTF genome annotation file obtained from Ensembl (release-102), using the command ‘cellranger-arc mkref’ (v2.0.0, 10x Genomics). Raw sequencing files were aligned to this custom reference genome and quantified using the command ‘cellranger-arc count’. Similar downstream processing was performed as described in the macaque multiome processing, expepte for some parameter changes (2000 variable genes were used for principal component analysis; top 20 principal components for transcriptome modality and top 16 singular values for chromatin accessibility modality were used for UMAP visualization; Louvain clustering resolution was set to 1).

### Cell type alignment across species

To match the cell type annotation during neocortex development across species, an integration and label transfer strategy was applied. First, gene names in the macaque and mouse multiome transcriptome modality were converted to their human homologous genes, using the ‘biomaRt’ R package (v2.54.0, Ensembl release version 105). Gene expression matrices in the macaque and mouse multiome data, along with the published human scRNA-seq data (Trevino, A.E. et al., 2021), were integrated using canonical correlation analysis (CCA) implemented in ‘Seurat’. Specifically, we ranked the 2000 variable genes in each dataset and selected 2000 shared genes for CCA reduction, based on their ranking. Top 30 dimensions in CCA reduction were used for searching the Mutual Nearest Neighbor (MNN) between datasets, which is also known as the anchors. Cell type annotation was transferred from the macaque multiome transcriptome modality to the mouse multiome transcriptome modality and the human scRNA-seq data, based on the anchor annotation around each cell in the query datasets.

To ensure the robustness of transcriptome integration, we explored additional parameters in the CCA method, including choosing top 40 and top 50 dimensions. We also incorporated the ‘harmony’ R package (v0.0.1) (*64*) for integration. Specifically, top 2000 variable genes shared among datasets were selected. Principle component analysis was then applied to the macaque multiome transcriptome modality based on these genes. Gene expression matrices from human and mouse were projected to the resulting PCA space. PCA embeddings from all datasets were then adjusted using the function ‘HarmonyMatrix’ (sigma parameter was set to 0.2 and theta parameter was set to 0.18). Label transfer and UMAP visualization were performed based on the harmonized PCA embedding.

### Data processing of the published scRNA-seq

Single-cell transcriptome sequencing is characterized by high dropout rate and noise, and there are technical differences between single-nucleus and single-cell transcriptome sequencing. In order to avoid drawing incorrect conclusions caused by these biases, we collected additional neocortex single-cell transcriptome data to validate our results, including one human scRNA-seq data from gestation week 17 to 18 (*13*) and one mouse scRNA-seq data from embryonic day 14 to 16 (*16*). By employing the aforementioned approaches, gene symbol annotations and cell type annotations were harmonized across datasets.

### Predictions of accessible genomic elements and gene expression linkage across cell types

Gene regulatory elements (GREs) are DNA sequences that modulate the transcription of target genes, playing a fundamental role in regulating gene expression. These elements, such as enhancers and promoters, primarily exert their effects on the genome in the form of open chromatin. Therefore, integration of chromatin accessibility and transcriptome data can aid in the inference of potential regulatory relationships between accessible genomic elements and gene expression.

In the analysis of multiome data from macaque and mouse, the ‘addPeak2GeneLinks’ function bulit in ‘ArchR’ was employed to compute the correlation between accessible elements and target genes using paired gene expression and peak accessibility matrices. Linkages with a correlation coefficient below 0.75 and distance exceeding 250,000 base pairs were excluded from the analysis. Consequently, 49,449 and 55,007 peak-to-gene linkages were identified in macaque and mouse, respectively. For scRNA-seq and snATAC-seq data from human, a constrained integration was initially executed using a similar approach as described in macaque snATAC-seq processing. Following the integration, pseudo-scRNA-seq profiles for each snATAC-seq cell were generated and employed for the peak-to-gene linkage calculation. Overall, 55759 linkages were identified across all cell types.

### Aggregated peak analysis

To evaluate the enrichment of peak-to-gene links identified by scMultiome data, the aggregated peak analysis (APA) was performed using the Hi-C profiles for the macaque cortex (*14*). Only peak-to-gene links with a distance between peak and gene that fell within the range of 50-250 kb were retained for analysis. The enrichment submatrices (observed counts divided by expected counts) around these links were summed. The Knight-Ruiz normalized Hi-C matrix with 5 kb resolution was used.

### Cortex laminae enrichment analysis

To identify cortex laminae-specific markers, differential expression analysis was performed on the macaque Laser Microdissection (LMD) bulk RNA-seq using ‘DESeq2’ (v1.38.3) (*60*). To assess the enrichment of these laminae-specific marker programs in the single-cell/single-nucleus sequencing data, we calculated the average expression levels using the ‘AddModuleScore’ function implemented in Seurat.

### Identification of cell markers per species

To enhance the accuracy and reliability of scRNA-seq and snRNA-seq, a two-step strategy was employed for cell marker detection. This strategy comprised identifying cell type-specific genes from a single-cell/single-nucleus transcriptome dataset, followed by cross-validation of these findings using an additional single-cell/single-nucleus transcriptome dataset.

In the first step, the gene expression percentage (Pct) and average expression level (Exp) were calculated for each cell type using the human scRNA-seq data, the macaque scMultiome data, and the mouse scMultiome data. The specificity of gene expression was evaluated by calculating two metrics: expression percentage specificity (Pct.diff) and expression level specificity (Exp.diff).

Expression percentage specificity was defined as:

Pct.diff = Pct[target cell type] – max(Pct[non-target cell types]) Expression level specificity was defined as:

Exp.diff = Exp[target cell type] / max(Exp[non-target cell types])

Genes with Pct[target cell type] surpassing 0.3, Pct.diff exceeding 0.2, and Exp.diff greater than 1 were retained.

In the second step, the gene expression percentage and average expression level were again calculated using additional transcriptome profiles. The human scRNA-seq data were obtained from (*13*); macaque data were procured from snRNA-seq, and mouse data were extracted from (*16*).

Genes identified in the first step with Pct[target cell type] surpassing 0.3, Pct.diff exceeding 0.1, and Exp.diff greater than 1 in the second step were recognized as consistent cell markers.

### Cross-species comparison of cell markers

In order to account for the high drop-out rate inherent in single-cell transcriptome data, a stringent threshold was employed to identify species-specific markers. Specifically, only markers exhibiting a significant difference in expression specificity between the target species and both non-target species were defined as species-specific markers. The expression specificity in macaque and mouse cell populations was evaluated based on the expression percentage, the average expression level, and the expression correlation with the human data.

For instance, in identifying the human-specific RG-neu markers, the human RG-neu markers with Pct.diff exceeding 0.1, Pct[target cell type] surpassing 0.15, Exp.diff greater than 1, and expression correlation greater than 0.9 in macaque or mouse were filtered as the macaque-shared or the mouse-shared markers.

### Gene Ontology (GO) analysis of the RG-neu signature genes

To elucidate the putative molecular/biological functions and cellular components attributed to the RG-neu signature genes across diverse species, Gene Ontology (GO) analysis was conducted by utilizing the ‘topGO’ R package (v2.50.0). The background gene set comprised all detected genes in the human scRNA-seq. The ‘classic’ algorithm in conjunction with the ‘Fisher’ statistical test was employed to evaluate the overrepresentation of GO terms sourced from ‘org.Hs.eg.db’ R package (v3.16.0).

### Genome accessibility comparison across species

To compare chromatin accessibility differences among genomic elements from various species, the coordinates and identities of these accessible regions must be harmonized. To achieve this, the ‘UCSC liftOver’ tool was utilized to convert accessible regions (peaks) from the macaque Mmul_10 and the mouse GRCm38 reference genomes to the human GRCh38 reference genome. The minimum ratio of bases required to remap was set at 0.9 for macaque and 0.6 for mouse. To further control for substantial size alterations caused by large rearrangements, deletions, or insertions, peaks exhibiting length change ratios exceeding 0.1 in macaque and 0.4 in mouse after liftover were excluded. Furthermore, peaks that either overlapped with one another or fell into unannotated regions following the liftover process were also removed. Peaks originated from human, macaque and mouse snATAC-seq data were then merged together to create a consensus peak set using ‘bedtools merge’ (v2.30.0). To ensure a one-to-one correspondence of this consensus peak set across species, the peak set was re-lifted over to the macaque and mouse genomes, adhering to the aforementioned filtering criteria. This step ensured that the final consensus peak set maintained a consistent representation of chromatin accessibility regions across the human, macaque, and mouse genomes. The accessibility score of this consensus peak set was represented by the corresponding genome’s Tn5 insertion counts in each species.

The DESeq2 and Wilcox algorithms were utilized to analyze accessibility score discrepancies between species. For the DESeq2 approach, cells within each cell type were arbitrarily divided into three pseudo-bulk samples. The Tn5 insertion counts were then combined within each pseudo-bulk sample and incorporated into ‘DESeq2’ (v1.38.3) to compute log2 fold change and p-values. In the case of the Wilcox method, insertions were initially normalized by the total insertion counts in peaks for each cell, followed by the calculation of log2 fold change and p-values. For both methods, the human-specifically gained chromatin accessible regions (HGARs) were identified as consensus peaks that exhibit a log2 fold change greater than 1 and a p-value below 0.01 when compared to macaque and mouse. These regions were also required to overlap with peaks detected in the human snATAC-seq data. The primate-conserved chromatin accessible regions (PCARs) were established as consensus peaks demonstrating a log2 fold change over 1 and a p-value less than 0.01 in comparisons of human and macaque to mouse, as well as an absolute log2 fold change between human and macaque below 1. Additionally, these regions had to overlap with human and macaque peaks in their respective snATAC-seq data. Conserved chromatin accessible regions (CARs) were defined as consensus peaks displaying an absolute log2 fold change of less than 1 when comparing human, macaque, and mouse pairwise. Moreover, these regions were required to overlap with peaks in the snATAC-seq data for each species.

The identified chromatin accessible regions were subsequently annotated for genomic distribution using the ‘ChIPseeker’ R package (v1.34.1). The gene annotation information was derived from the ‘org.Hs.eg.db’ R package and the Homo sapiens GRCh38 GTF file (Ensembl release 105). The transcription start site (TSS) region was defined as 2000 bp upstream and downstream of the TSS.

### Evolutionary origin of HGARs

To explore the evolutionary origins of HGARs across different cell types, HGARs were classified into four distinct categories: de novo, co-option, co-option PFC, and modification. This classification was based upon the accessibility profiles observed in the macaque prefrontal cortex (PFC), the mouse PFC, and the mouse non-PFC fetal tissues. For the purpose of comparison, accessible regions (ARs) from nine non-PFC mouse fetal tissues obtained from the ENCODE project were also utilized.

For instance, the de novo HGARs in the RG-neu cell types were defined as those that lacked any detected ARs in the macaque PFC, the mouse PFC, and the mouse non-PFC tissues. HGARs with detected ARs solely in the mouse non-PFC tissues were identified as co-option HGARs. Co-option PFC HGARs were classified as HGARs absent of detected ARs in RG-neu of both macaque and mouse PFC, while modification HGARs were categorized as HGARs with detected ARs in either RG-neu of macaque or mouse PFC.

### Human-fixed SNC and HAR enrichment analysis

The enrichment of human genome features in conserved and species-specific ARs was evaluated. The human-fixed SNC position file was downloaded from http://ftp.ensembl.org/pub/data_files/homo_sapiens/GRCh38/compara/. The HAR position file was sourced from previous studies (*26, 39, 40*). The density of the human-fixed SNC or HAR with ARs was calculated as the average number of SNCs or HARs overlapping with ARs per kilobase. The enrichment of HARs was quantified as the fold change in density relative to the density of all ARs. The statistical significance of the enrichment was assessed using Fisher’s exact test.

### Scanning for motif occurrence in HARs containing the human-fixed SNCs

The emergence of HGARs may be potentially attributed to the occurrence of specific nucleotide sequence alterations in the homologous genomic regions within the human lineage, which may introduce new transcription factor binding sites. Under the influence of transcription factor binding, these regions exhibit a human-specific open chromatin state. To investigate this hypothesis, the human-fixed SNC sites were extended by 30bp on each end. Sequences that intersected with HGARs were retained and introduced to FIMO for motif site scanning. The motif file was sourced from JASPAR2022. Among the motif binding site scanning results returned by FIMO, only those with a p-value less than 10^-4 and intersecting with the human-fixed SNC were retained.

### Linkage disequilibrium score regression (LDSC) analysis

LDSC was implemented to investigate the enrichment of HGARs over disease-associated SNPs. All HGARs were converted from GRCh38 to GRCh37. The LDSC analysis was conducted in accordance with the tutorial (https://github.com/bulik/ldsc/) as previously described (*65, 66*). GWAS summary statistics for intelligence, schizophrenia, neuroticism, bipolar disorder, autism spectrum disorder, attention-deficit hyperactivity disorder, Parkinson’s disease, Alzheimer’s disease, anxiety, anorexia, and epilepsy were utilized. The trait for body mass was employed as a control.

### RNA Fluorescent in situ hybridization (RNA FISH)

The FISH experiments were performed using the RNAscope Multiplex Fluorescent Reagent (ACD Bio) as described. Brains were sliced in 15 μm thick with coronal sections. The *ITGA2*, *GPX3*, *PAX6* and *FOS* RNAscope Probes were designed and RNAscope® Multiplex Fluorescent Reagent Kit v2 were used to detect these transcripts.

### Quantification of RNA FISH

In order to ascertain the expression pattern of the human-specific RG-neu marker gene across species, RNAscope images were analyzed using ImageJ version 1.53t. For each image, regions of interest (ROIs) were manually outlined in the VZ layer (human, macaque, and mouse) and OSVZ layer (human and macaque), based on the nuclear packing determined by DAPI staining. For each ROI, the integrated fluorescence intensity was separately calculated for the red, green, and blue channels. To rectify the variations in cell density among different ROIs, the integrated fluorescence intensity for the red and green probes was divided by the integrated DAPI fluorescence intensity from the blue channel, resulting in the normalized fluorescence intensity. In total, each probe obtained between 67 and 167 valid data points for each species.

### In utero electroporation(IUE)

Pregnant ICR mice (E13.5) were used for in utero electroporation. The human *ITGA2* gene was cloned into the PCAGGS plasmid vector, and the empty PCAGGS plasmid vectors were used as the control. IUE was performed as previously described (*67*). In brief, plasmids were mixed with 0.1% Fast Green and the injection concentration was adjusted to 1.5 μg/μL. About 1 μL plasmid solutions were injected into the lateral ventricles of embryonic brain. Mice were continuously anesthetized with isoflurane during surgery. Electroporation was performed using BTX830 electroporator according to the following conditions (five pulses of 35V for 0.05s with the pulse interval of 0.95s). After electroporation, the embryos were returned to the abdominal cavity with suitable warm saline, and mice were sutured and placed on room temperature until recovery. The embryonic brains were collected in E15.5 and E18.5 after the pregnant mice sacrificing and analyzed at least four different brains for each group.

### Immunofluorescent staining

Brains were fixed in 4% PFA for 12h, and placed at 4 °C overnight to allow tissues to sink into 15%-30% gradient sucrose solution. Brains were sliced in 13 μm thickness in preparation. For immunohistochemistry, brain sections were placed in antigen retrival solution (Servicebio, G1206) and steamed for 10mins in 90℃. Sections were subsequently permeabilized with PBST (1x PBS+0.2% Triton X-100 (Sigma-Aldrich, T8787)) for 30 mins, and blocked with 5% BSA for 1h in 37 ℃. Then, brain sections were incubated with diluted primary antibody overnight at 4°C and diluted secondary antibody for 1h at 37°C, followed by DAPI (Thermo Fisher Scientific,62248, 1:1000) staining for 1 min at room temperature. Brain sections were mounted by ProLong® Gold Antifade reagent (ThermoFisher Scientific, P36930). The following primary antibodies were used: GFP (abcam, ab13970, 1:1000), PAX6 (BioLegend, PRB-278P, 1:500), TBR2 (abcam, ab23345, 1:500), KI67 (abcam, ab16667, 1:500), TBR1 (abcam, ab31940, 1:500), SATB2 (abcam, ab51502, 1:200), CRYAB (abcam, ab13496, 1:500), ANXA1(abcam, ab214486, 1:500), AlexaFluor488 Goat anti-chicken (abcam, ab150169, 1:1000). Secondary antibodies: AlexaFluor594 Donkey anti-rabbit (abcam, ab150076, 1:1000), AlexaFluor647 Donkey anti-mouse (Invitrogen, A-31571, 1:1000). Digital image acquisition was performed using LEICA TCS SP8 microscope.

### Quantification of immunofluorescent staining

IUE embryonic brains with at least four biological replicates per time point were included in the analysis, and only sample pairs with the similar electroporation efficiency and comparable brain regions between treatment and control were kept for further analysis. Each brain was selected three brain sections as the technical replicates. Using the annotation function of Leica confocal microscope, each brain section was delineated 100-μm wide for counting.

### Data and materials availability

The snMulitome data for macaque and mouse PFC, snRNA-seq, scATAC-seq, LMD bulk RNA-seq and bulk ATAC-seq data were deposited at NCBI GEO under the accession number: GSE241429. Scripts used in this study are available at Github repository: https://github.com/yimingsun12138/BrainEvoDevo. All data are available in the main text or the supplementary materials.

## Supplemental figures

**Fig. S1.**
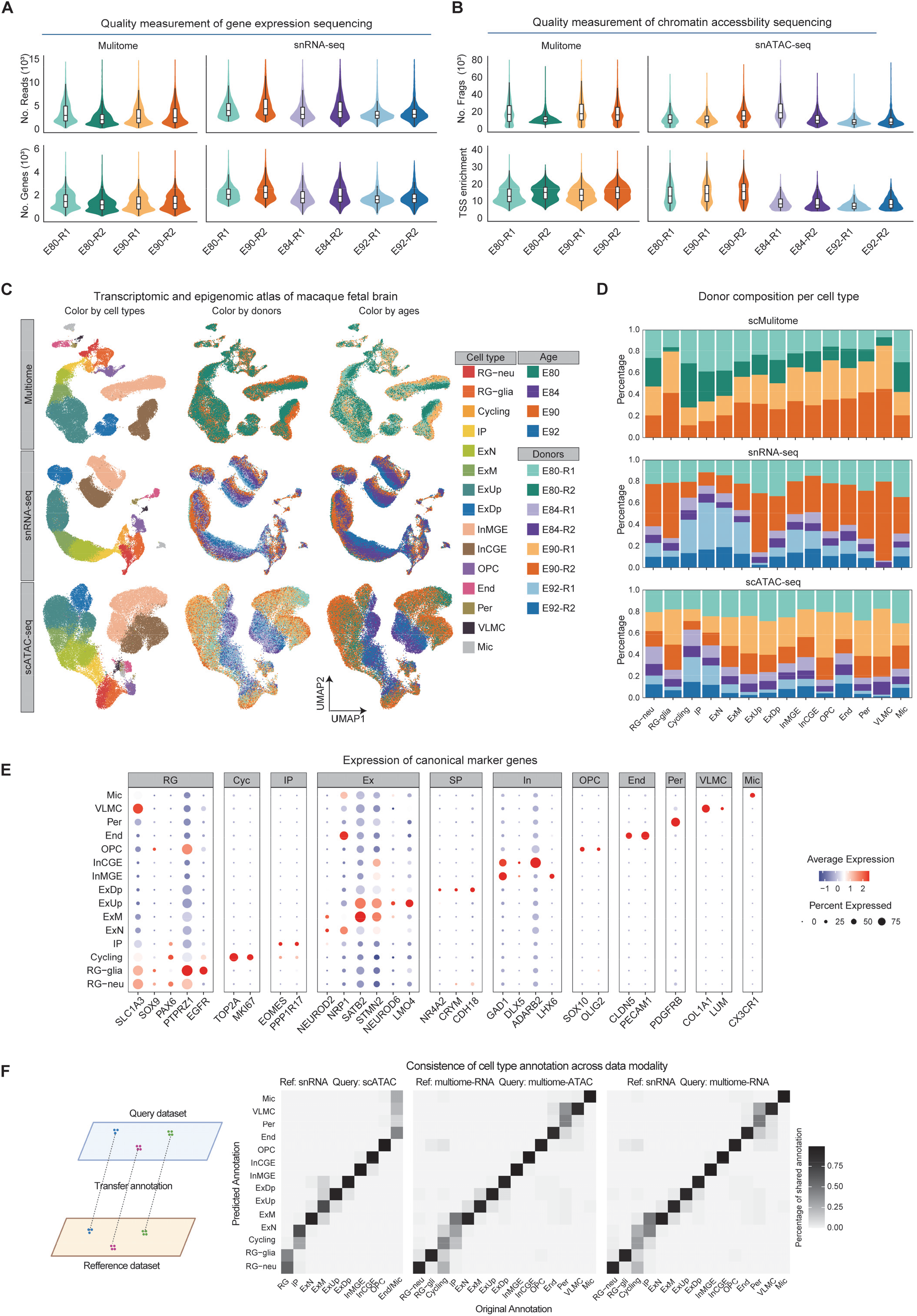
Cell atlas constructed from the single-nucleus transcriptome of the macaque PFC during corticogenesis. **(A)** Number of transcripts (top) and genes (bottom) detected for each single cell that passed the filter in each experimental sample. E, embryonic day; R, replication index for each sample. **(B)** Number of nuclear sequencing fragments (top) and normalized enrichment score (bottom) of the sequencing fragments around the transcription start sites (TSS) for each single cell that passed the filter in each experimental sample. **(C)** Overlay of cell types, biological replicates, and development time labels on a UMAP visualization of macaque fetal neocortex single-nucleus RNA sequencing (snRNA-seq), single-nucleus Assay for Transposase-Accessible Chromatin sequencing (snATAC-seq), and single-cell Multiome data (scMultiome). **(D)** Bar plot illustrating the distribution of composition for each cell type of the biological replicates in the macaque fetal neocortex snRNA-seq, snATAC-seq and scMultiome data. **(E)** Bubble plot of expression profiles of canonical cell-type marker genes. **(F)** Left panel shows the prediction of cell type labels in the reduced-dimensional space based on the proximity relationship between cells (see materials and methods). Right panel illustrates the shared percentage of the original annotation labels and the predicted annotation labels under three distinct scenarios: predicting snATAC-seq cell type labels using snRNA-seq data; predicting snATAC-seq cell type labels in sscMultiome data utilizing the transcriptome data from the same scMultiome data; predicting transcriptome cell type labels in scMultiome data employing snRNA-seq data.

**Fig. S2.**
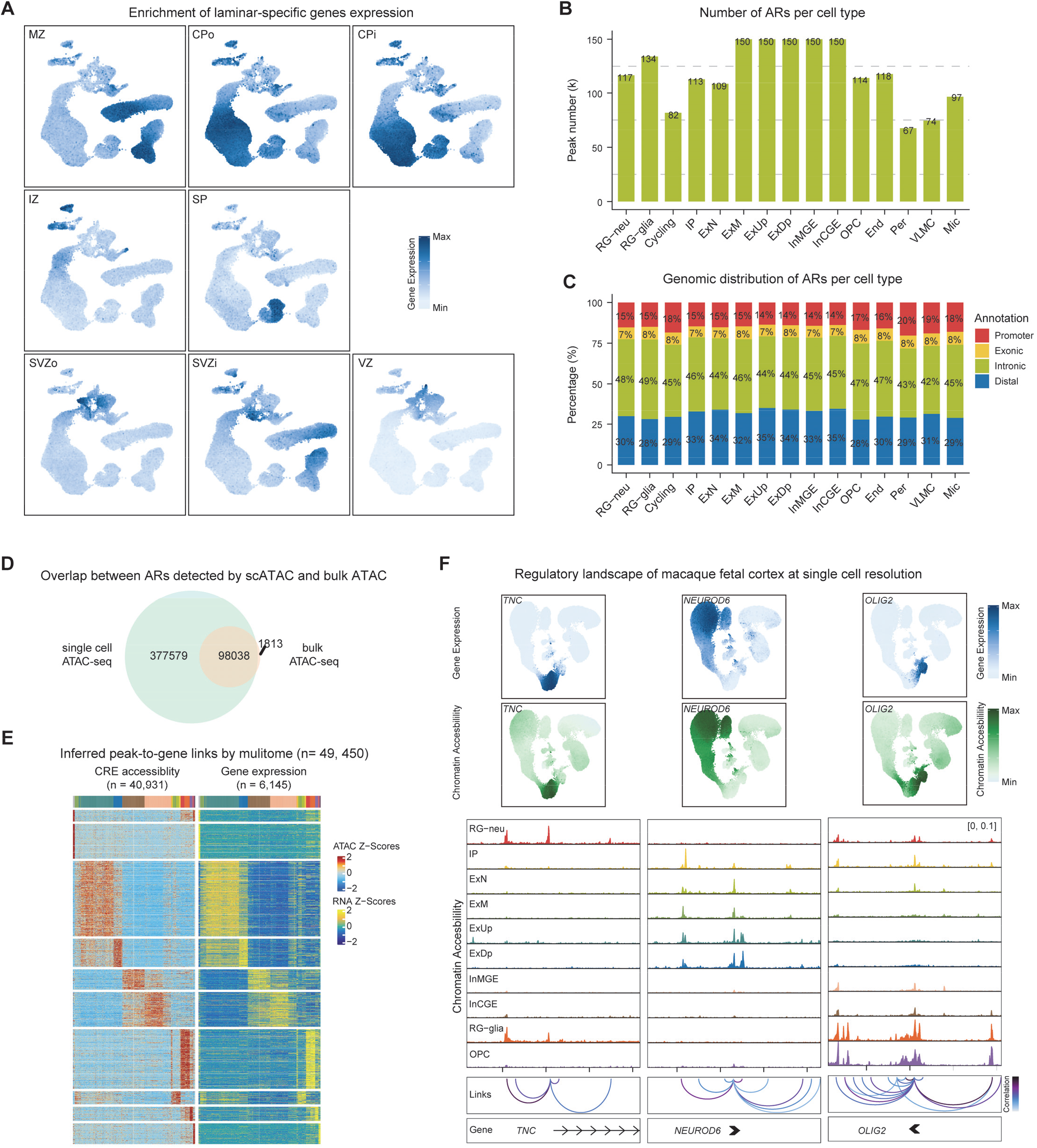
Cell atlas based on the regulatome of the macaque PFC during corticogenesis. **(A)** UMAP plot of the macaque scMultiome data illustrates the expression module score of the fetal neocortex laminar-specific genes identified using the macaque LMD bulk RNA-seq. **(B)** Bar plot illustrates the quantity of the detected open chromatin peaks, also known as accessible regions (ARs), for each cell type annotated in the macaque scMultiome data. **(C)** Distribution ratio of the open chromatin peak annotations for each cell type in the macaque scMultiome data. **(D)** Venn diagram illustrates the intersection of the open chromatin peaks detected using both the macaque bulk ATAC-seq data and the scMultiome data. **(E)** Heatmap displays the correlation between the scaled peak accessibility (left) and gene expression (right). Each row represents a peak-to-gene link detected by ArchR, while each column corresponds to a meta cell in scMultiome data, individually matching the transcriptome assay and chromatin accessibility assay. **(F)** Scaled gene expression (top) of the canonical marker genes for neuron progenitor cells (TNC), excitatory neurons (NEUROD6), and glial cells (OLIG2), along with the corresponding chromatin accessibility landscape and the predicted peak-to-gene linkage (bottom) surrounding these genes.

**Fig. S3.**
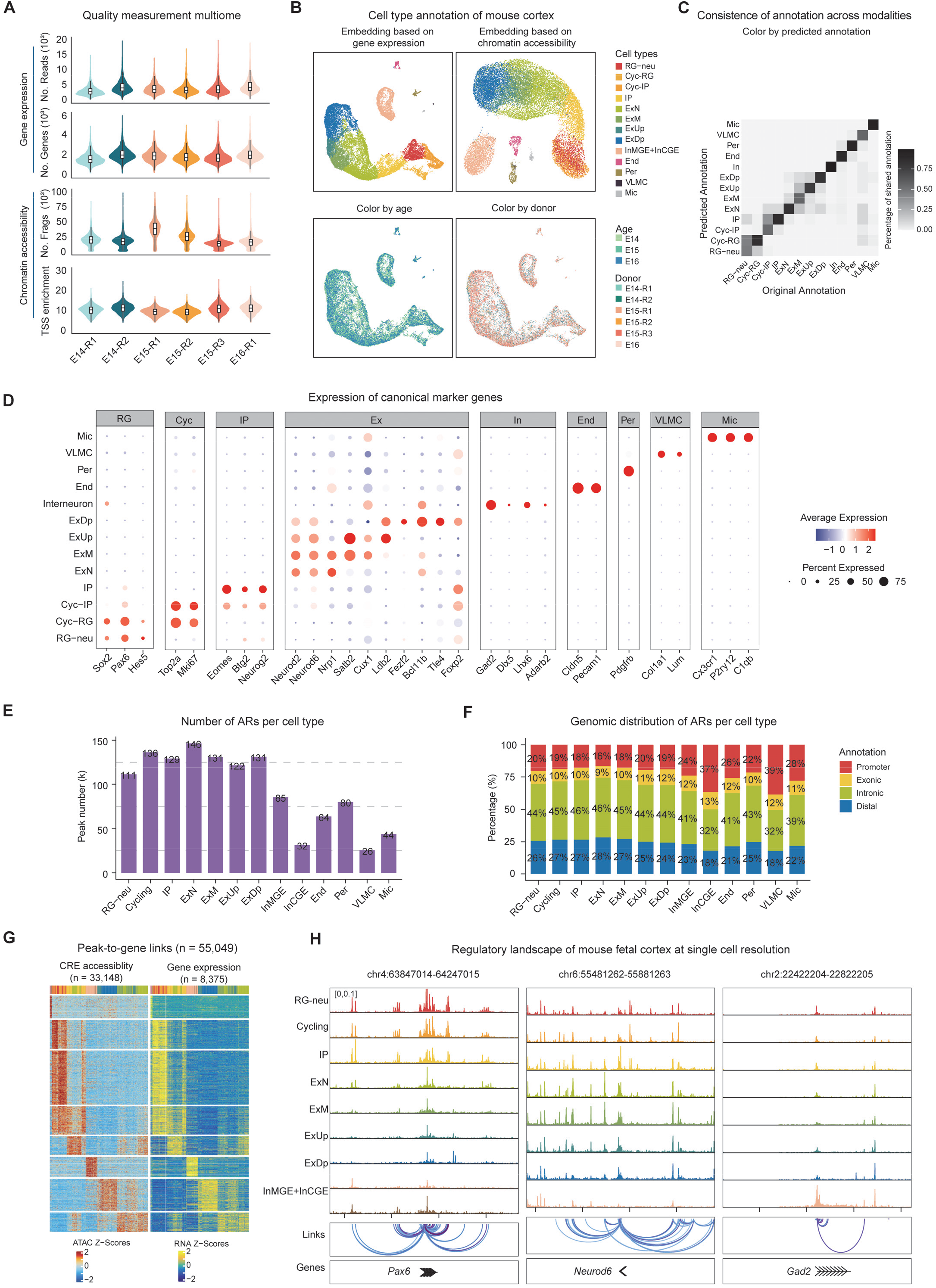
Cell atlas based on the transcriptome and regulatome of the mouse PFC during corticogenesis. **(A)** Quality control measurements of transcriptome (top) and chromatin accessibility (bottom) assays in the mouse scMultiome data. **(B)** UMAP charts of the mouse scMultiome transcriptome assay and chromatin accessibility assay, overlaid respectively with labels for cell type, development time, and biological replicates. **(C)** Heatmap illustrates the percentage overlap between the original annotation labels from the mouse scMultiome chromatin accessibility assay and the predicted annotation labels derived from the transcriptome assay. **(D)** Bubble plot of expression profiles of the canonical cell-type marker genes. **(E)** Bar plot illustrates the quantity of the detected open chromatin peaks for each cell type. **(F)** Distribution ratio of the open chromatin peak annotations for each cell type. **(G)** Heatmap displays the correlation between the scaled peak accessibility (left) and gene expression (right). **(H)** Chromatin accessibility landscape and predicted peak-to-gene linkage surrounding canonical marker genes for neuron progenitor cells (Pax6), excitatory neurons (Neurod6), and inhibitory neurons (Gad2).

**Fig. S4.**
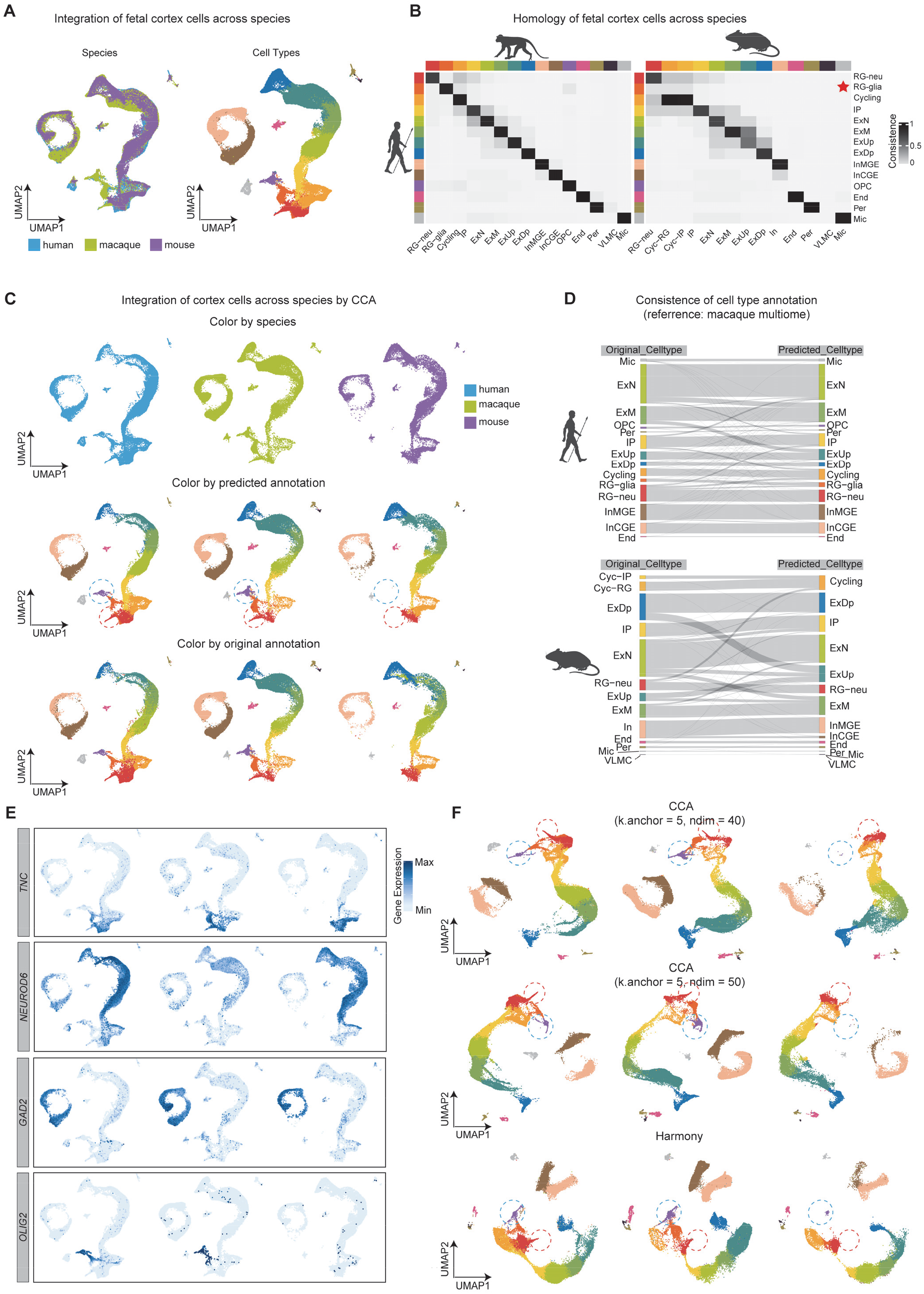
Integration of the transcriptomic profiles across species. **(A)** Harmonized UMAP embedding of human, macaque, and mouse transcriptome assays, generated through the CCA method of Seurat. **(B)** Heatmap illustrates the homology scores between various cell types across different species. **(C)** Harmonized UMAP plot that features an overlay of species labels, the original cell type annotations for each species, and the predicted cell type labels based on the macaque data. **(D)** Sankey plot demonstrates the relationship between the originally annotated cell type labels in human (top) and mouse (bottom), and the predicted cell type labels derived from the macaque data. **(E)** Overlay of scaled gene expression of the canonical marker genes for neuron progenitor cells (TNC), excitatory neurons (NEUROD6), inhibitory neurons (GAD2), and glial cells (OLIG2) on species harmonized UMAP plot. **(F)** Harmonized UMAP plots of human, macaque, and mouse transcriptome assays generated using the CCA method with varying parameters (top and middle) and the Harmony method (bottom). Red and blue circles represent two cell types that are exclusively found in the human and macaque datasets, but not in the mouse dataset.

**Fig. S5.**
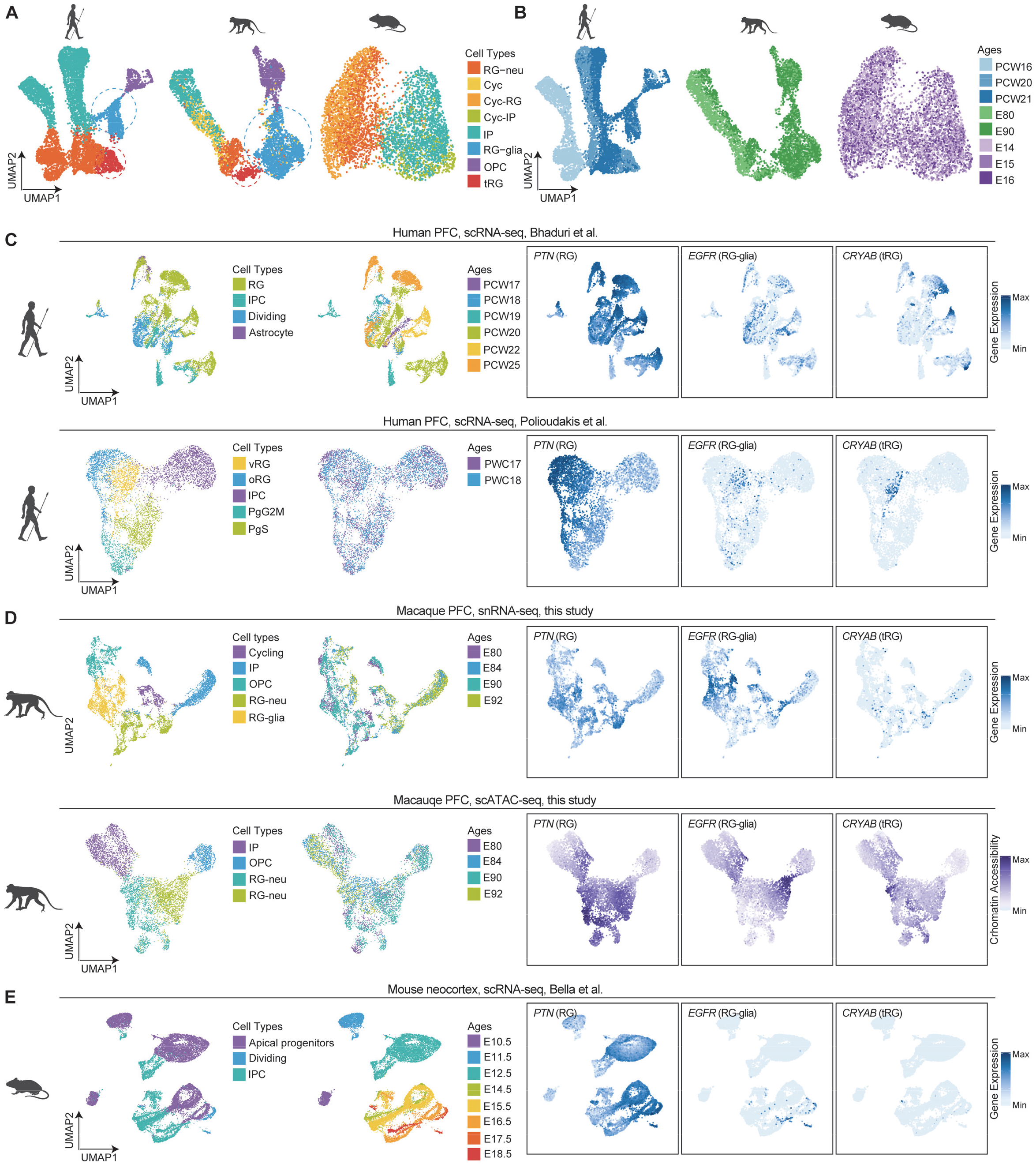
Validation for the primate-specific progenitors. **(A)** UMAP plots of neural progenitor cells from human (left), macaque (middle), and mouse (right), overlaid with the re-clustered and re-annotated cell type labels. **(B)** Overlay of fetal development time on the UMAP plots in (A). **(C)** Overlay of cell type annotation, fetal development time, and canonical marker gene expression on the UMAP plots of the human PFC scRNA-seq from (*68*) and (*13*). **(D)** Overlay of cell type annotation, fetal development time, and canonical marker gene expression on the UMAP plots of the macaque snRNA-seq and snATAC-seq. **(E)** overlay of cell type annotation, fetal development time, and canonical marker gene expression on the UMAP plots of the mouse PFC scRNA-seq from (*16*).

**Fig. S6.**
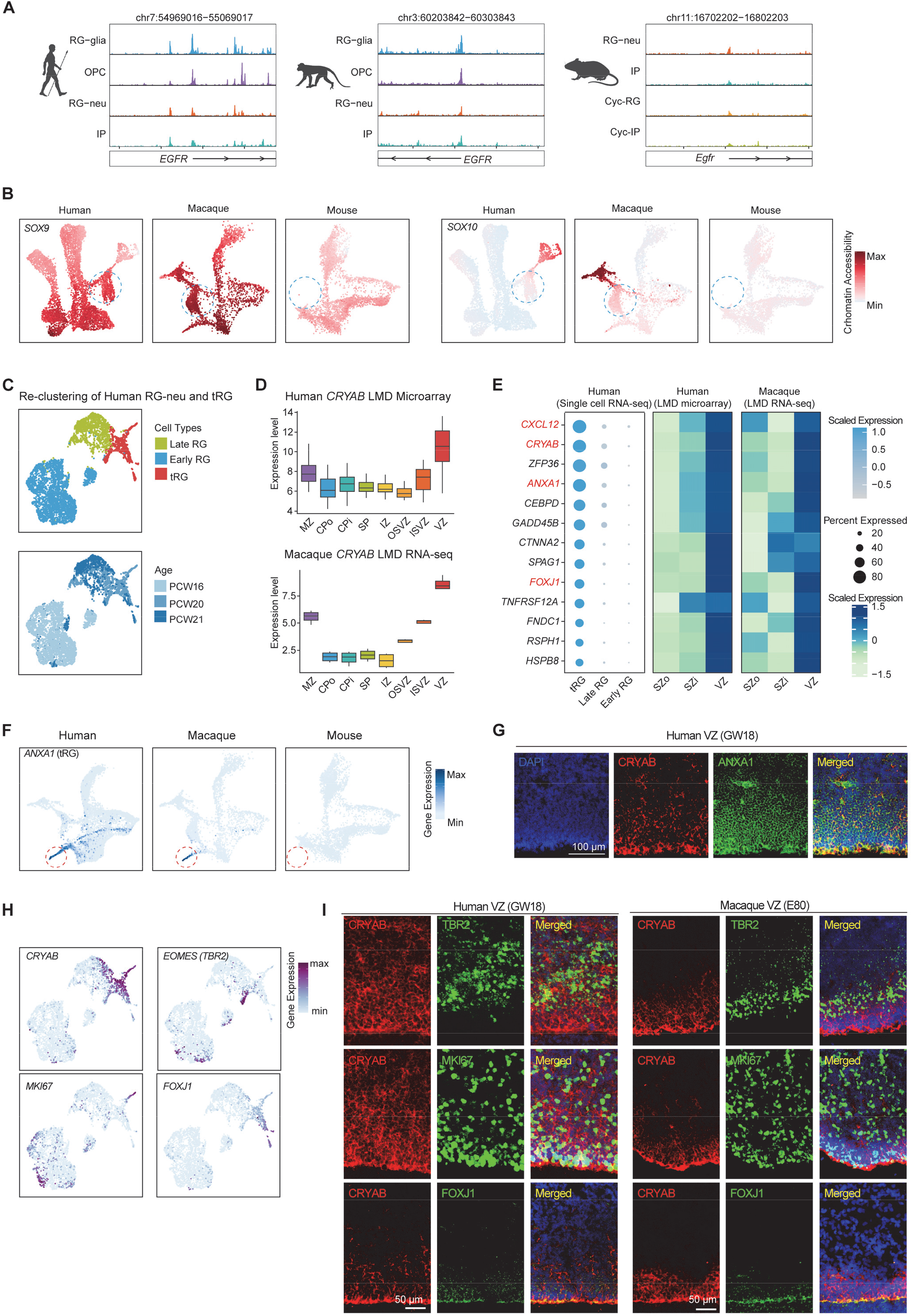
Molecular features of RG-glia and tRG. **(A)** Chromatin accessibility profiles around the classic RG-glia marker *EGFR* in three species. **(B)** UMAP plot illustrates the chromatin accessibility of *SOX9* and *SOX10* in progenitor cells across species. **(C)** UMAP visualization of all RG-neu in the human scRNA-seq dataset. **(D)** Laminar expression pattern of the canonical tRG marker *CRYAB* based on the LMD bulk RNA-seq. **(E)** Laminar expression pattern of the canonical tRG markers based on the LMD bulk RNA-seq. **(F)** UMAP plot illustrates the expression of the tRG marker *ANXA1* in progenitor cells across species. **(G)** Immunofluorescence staining for the tRG marker *ANXA1* in the human VZ. **(H)** Molecular features of the three branching directions in tRG differentiation. **(I)** Immunofluorescence staining for TBR2 (EOMES), MKI67 and FOXJ1 in the human and macaque VZ.

**Fig. S7.**
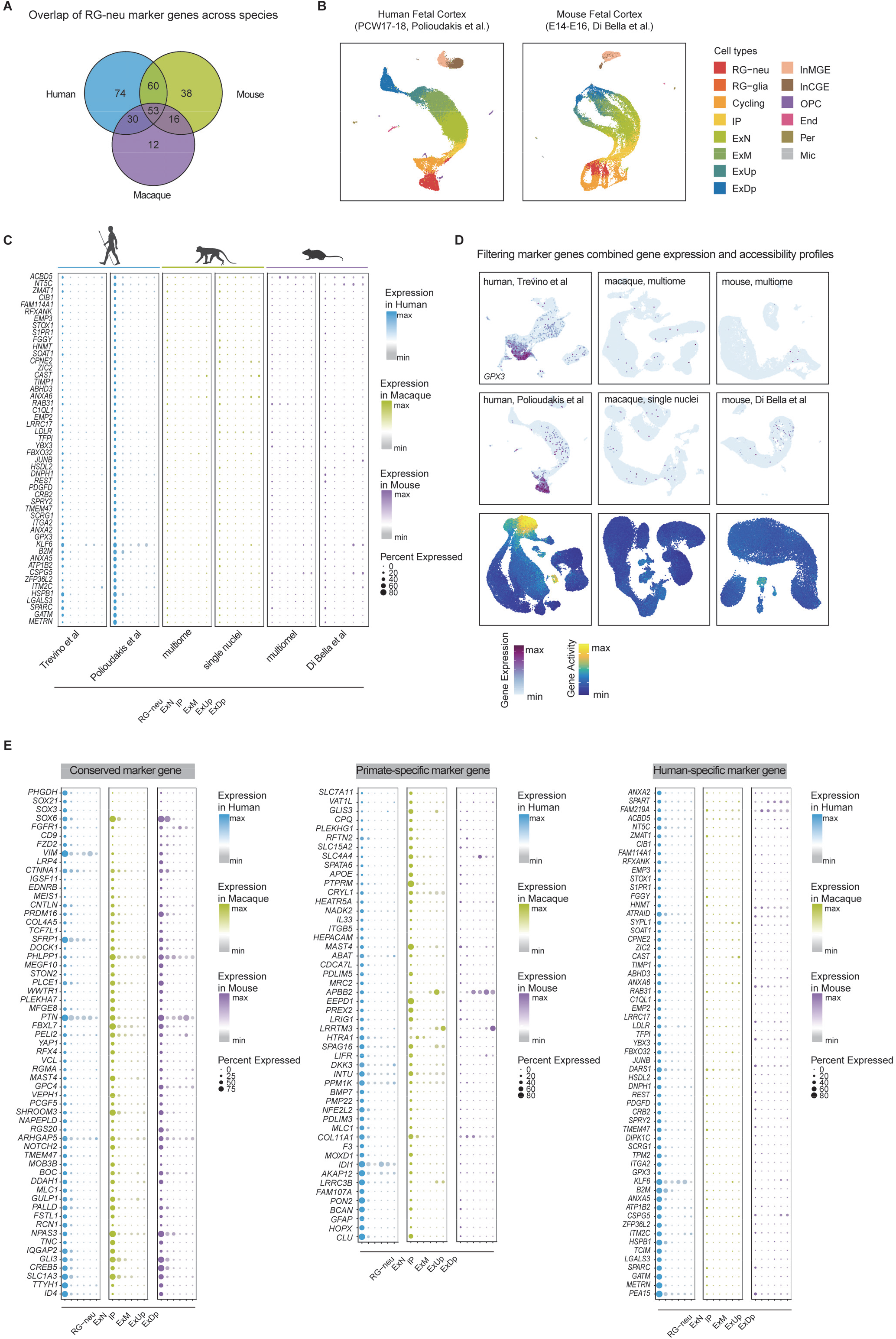
Cross-species transcriptomic comparison of the RG-neu markers. **(A)** Venn diagram illustrates the intersection of the RG-neu marker genes detected in each species. **(B)** UMAP plot incorporating two additional scRNA-seq datasets from the human (left) and mouse (right) fetal neocortex, utilized for the identification of consensus RG-neu markers. **(C)** Demonstration of the use of multiple scRNA-seq datasets for identification of datasets-conserved markers. **(D)** Demonstration of the use of gene expression and chromatin accessibility score for screening the human-specific RG-neu marker genes, using *GPX3* as an example (see materials and methods). **(E)** Bubble plots illustrates the expression of the conserved RG-neu markers (left), the primate-specific RG-neu markers (middle), and the human-specific RG-neu markers (right) across various cell types and species.

**Fig. S8.**
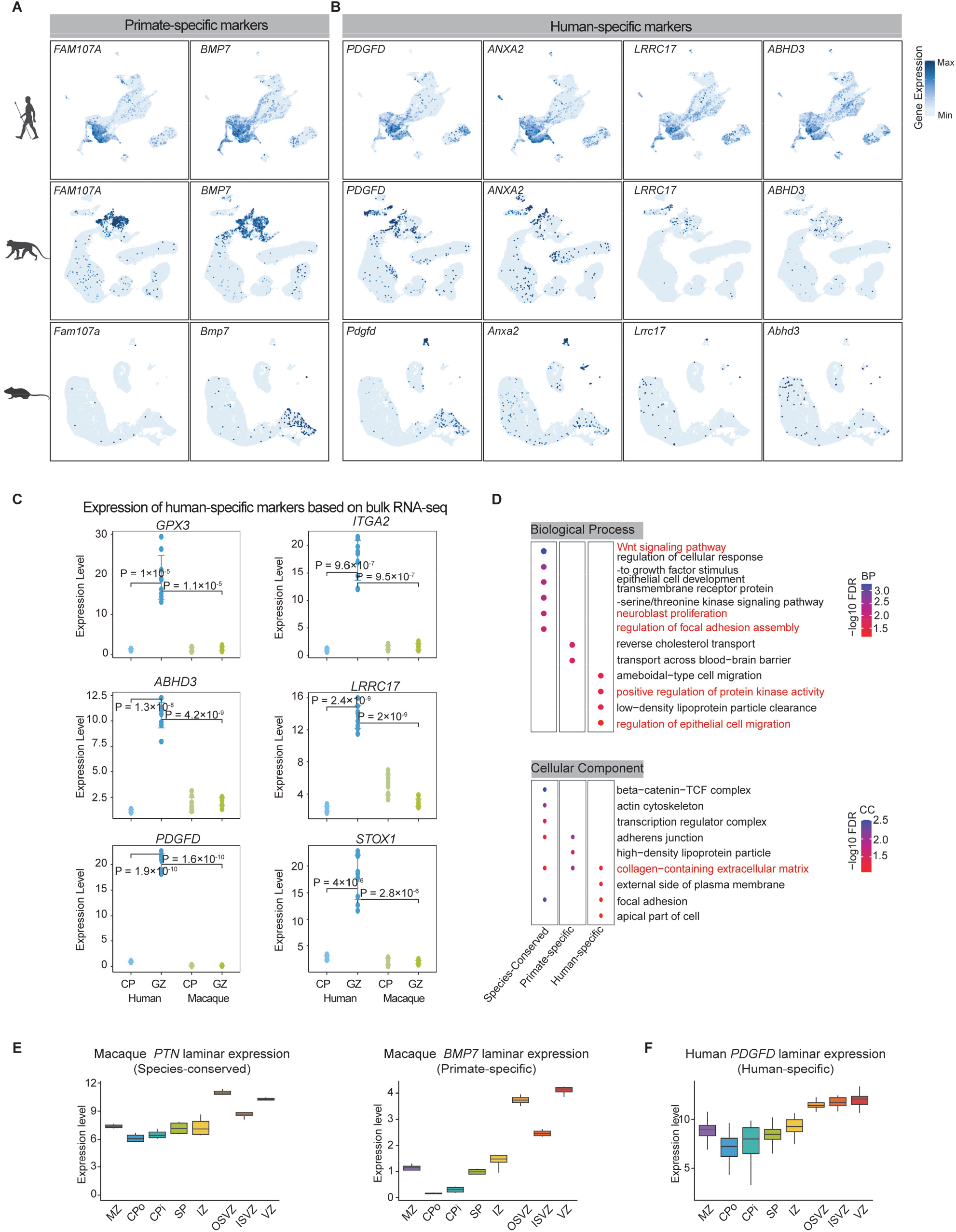
Validation and functional features of the primate-specific and the human-specific RG-neu markers. (A) UMAP plot illustrates the primate-specific RG-neu marker gene expression. (B) UMAP plot illustrates the human-specific RG-neu marker gene expression. (C) Validation of the human-specific RG-neu marker genes based on the bulk RNA-seq data. (D) Functional enrichment for the conserved and the species-specific RG-neu marker genes. (**E**-**F**) Expression patterns for the primate-specific (E) and the human-specific (F) RG-neu marker genes associated with growth factors.

**Fig. S9.**
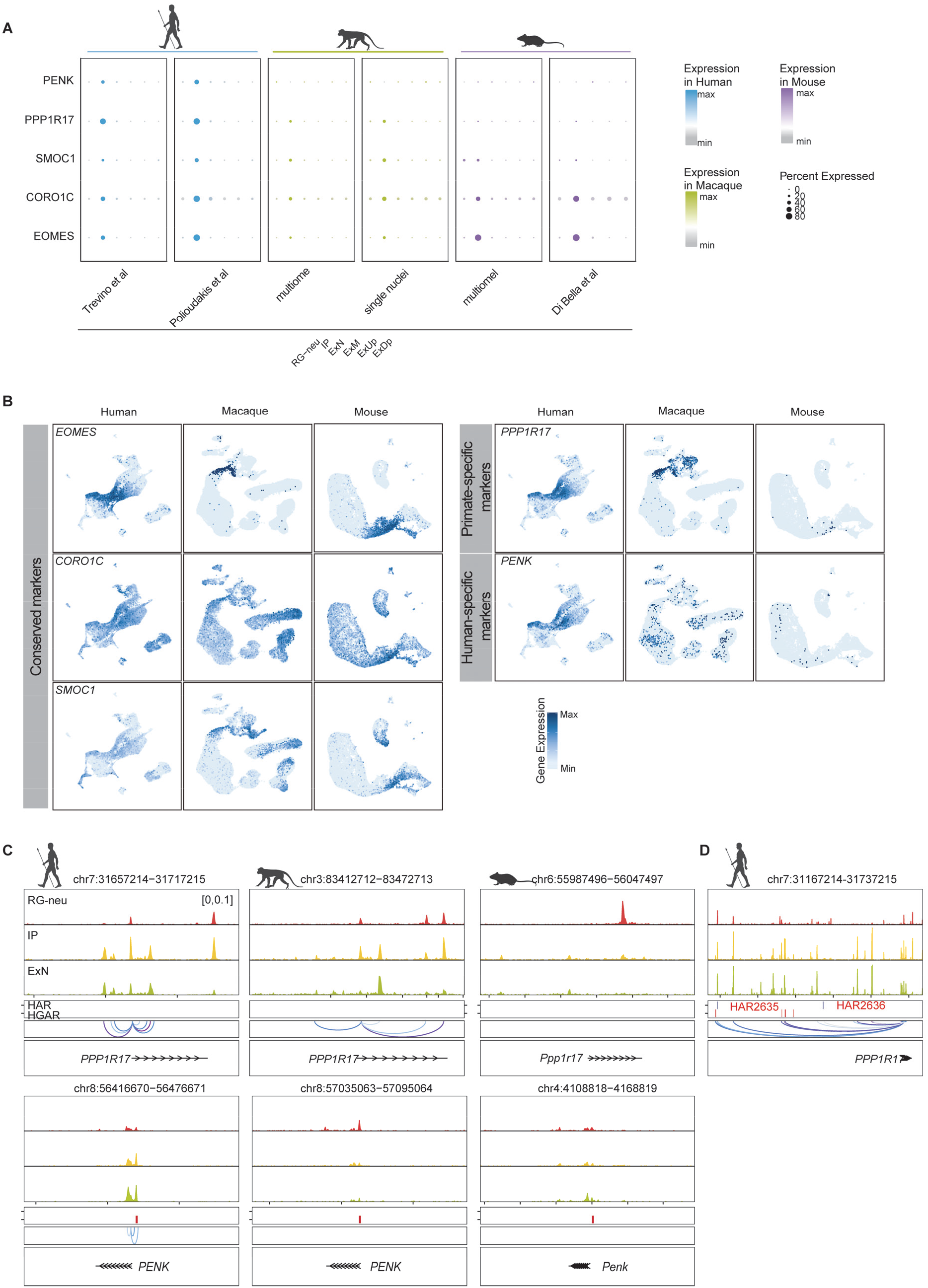
Species conservation for IPC markers. **(A)** Bubble plot illustrates the expression of the IPC marker genes in each species. **(B)** UMAP plot illustrates the expression of the conserved, the primate-specific, and the human-specific IPC marker genes in human, macaque and mouse. **(C)** Chromatin accessibility landscape and the predicted peak-to-gene linkages surrounding *PPP1R17* (top) and *PENK* (bottom). **(D)** Chromatin accessibility landscape surrounding the two human accelerated regions (HARs) predicted to be linked with *PPP1R17* in human.

**Fig. S10.**
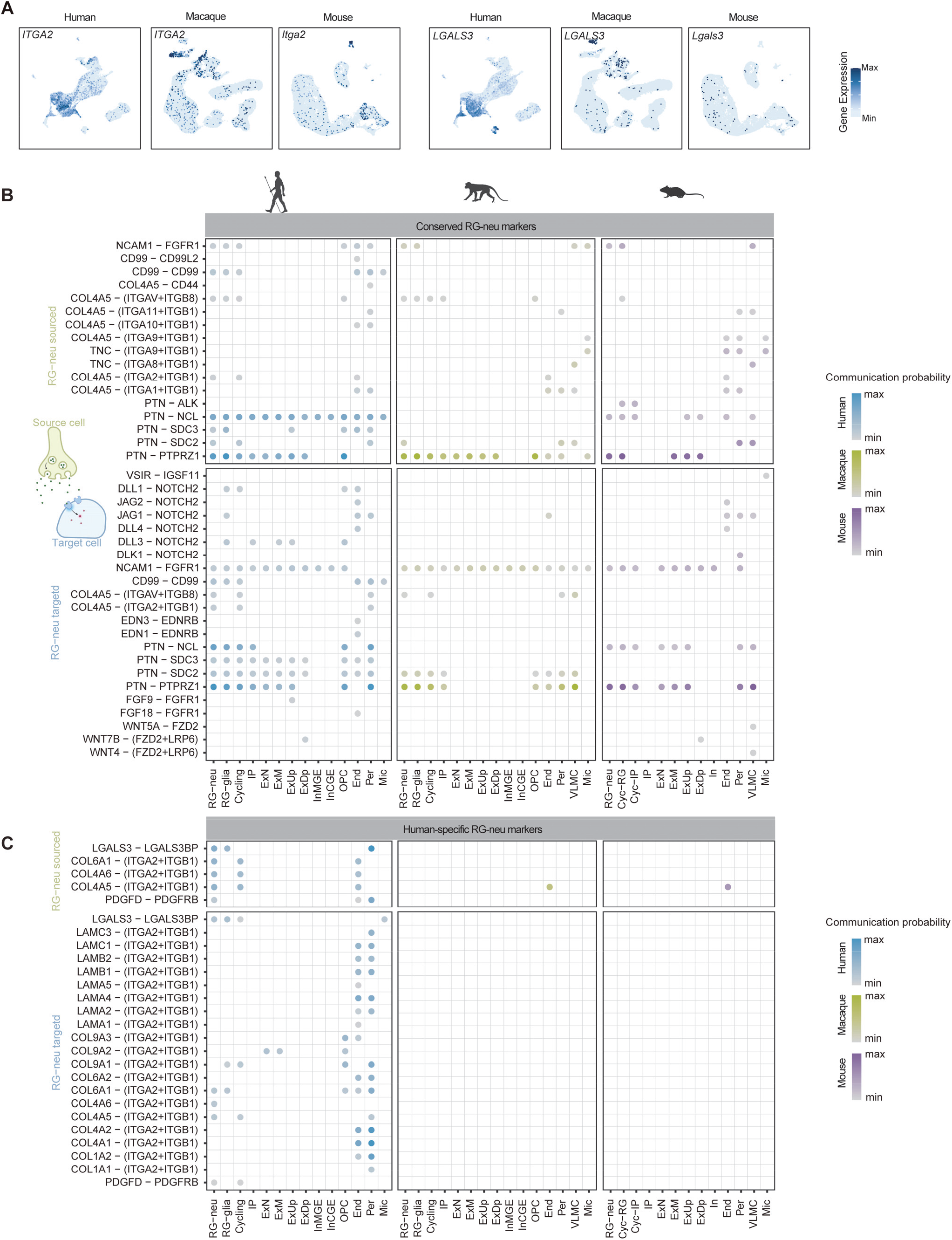
Human-specific cell communications between progenitor and blood cells. (A) UMAP plot illustrates the expression of *ITGA2* and *LGALS3* across species. (B) and **(C)** Cell communications mediated by the conserved (B) and the human-specific (C) RG-neu marker genes, where each row constitutes a set of ligand-receptor pairs, and each column represents a cell type, serving as the donor (top) and the receiver (bottom), respectively.

**Fig. S11.**
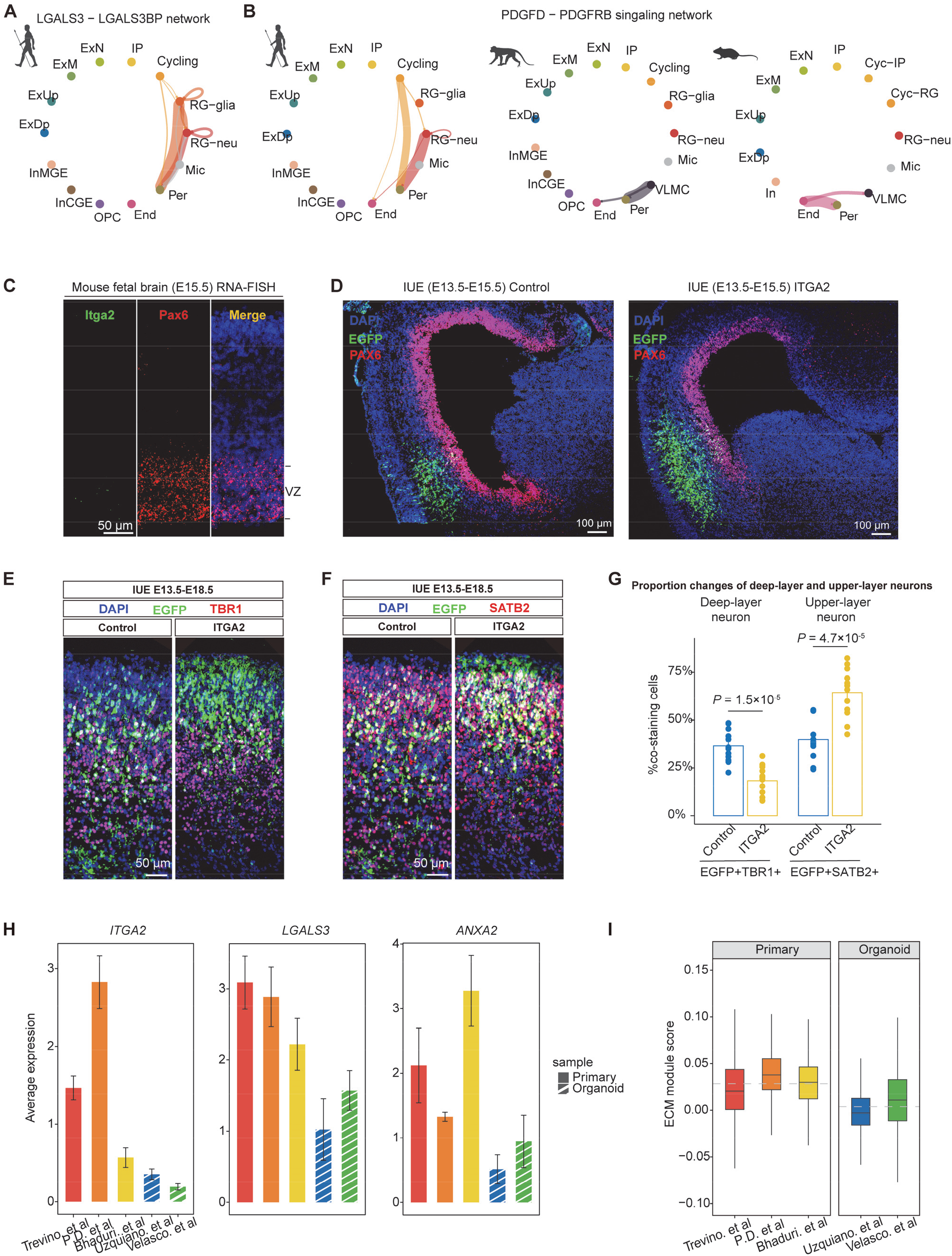
Functional implications of the human-specific RG-neu markers in corticogenesis. **(A)** and **(B)** The *LAGLS3* and *PDGFD* mediated cell-cell communication across species. (C) RNA FISH for the human-specific RG-neu marker *Itga2* in the mouse fetal cortex. (D) *In utero* electroporation of *ITGA2*+EGFP and EGFP alone (control) in the mouse cortex at E13.5 and analyzed at E15.5. (E) and (**F**) In utero electroporation of ITGA2+EGFP or EGFP only (negative control) in the mouse cortex at E13.5. The fetal cortex was analyzed at E18.5. Co-immunofluorescence-staining of ITGA2 with two neuron marker genes, including TBR1 for deep-layer neuron (E), and SATB2 for upper-layer neurons (F). Compared to the controls, there are significant increases of upper-layer neurons in the mouse fetal cortex subject to ITGA2+EGFP electroporation. (G) Bar plot illustrates the proportion changes in control and ITGA2+ electroporation mouse. P values were calculated by t test. (H) Bar plot illustrates the expression of *ITGA2*, *LGALS3* and *ANXA2* in the human primary cortex and the human brain organoids. **(I)** Box plot illustrates the ECM module score in the human primary cortex and the human brain organoids.

**Fig. S12.**
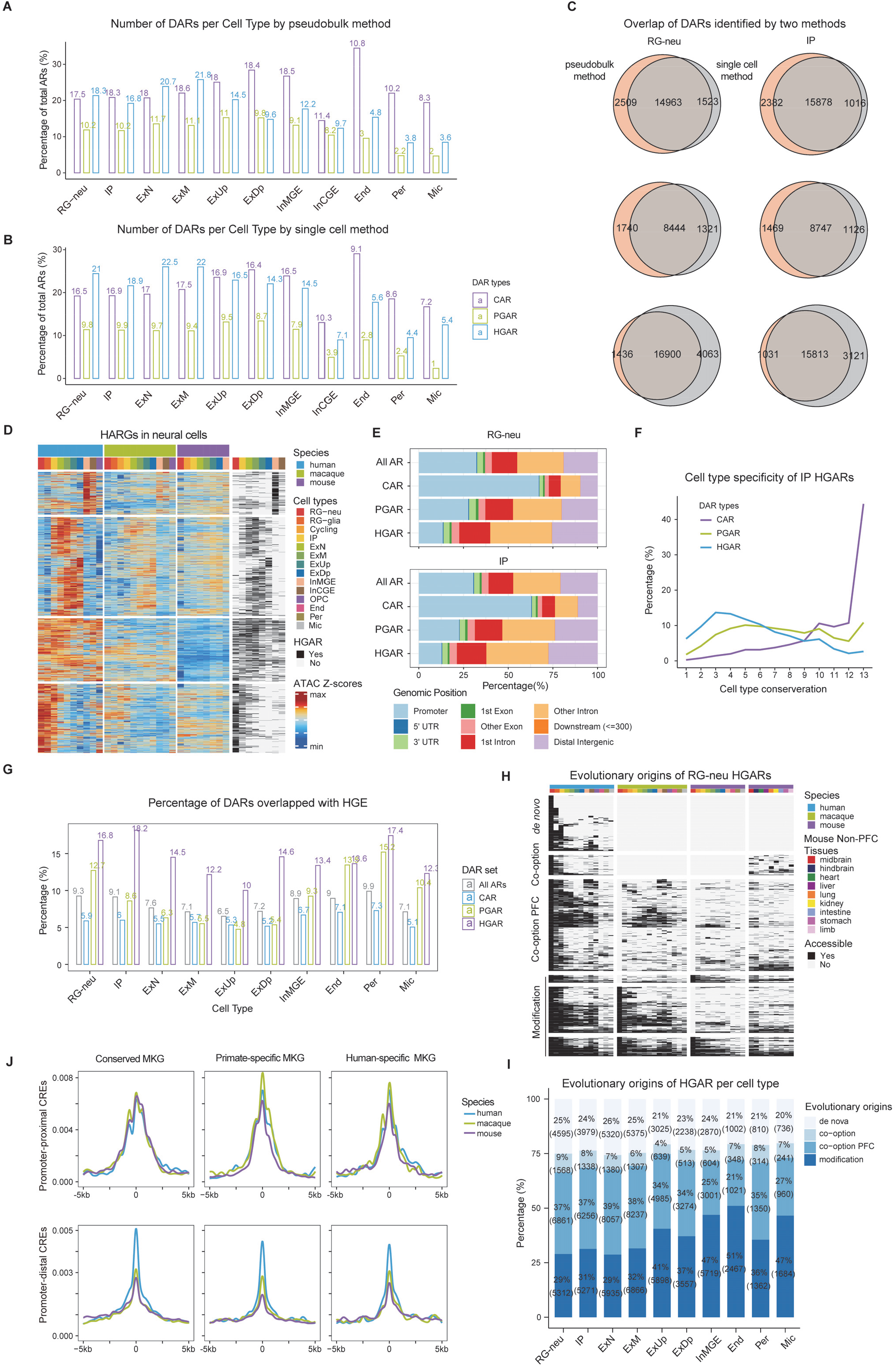
Identification and characterization of HGARs. **(A)** Bar plot illustrates the count of the conserved accessible region (CARs), the primate-gained accessible regions (PGARs), and the human-gained accessible regions (HGARs), calculated by DESeq2 in various cell types across species. **(B)** Bar plot illustrates the numbers of CARs, PGARs, and HGARs, calculated by the Wilcox algorithm in various cell types across species. **(C)** Venn diagram illustrates the overlap of differentially accessible regions (DARs) identified by both the DESeq2 and the Wilcox algorithms with various sampling ratios. **(D)** Heatmap illustrates the scaled counts of Tn5 insertions in HGARs across a range of cell types and species (displayed on the left). Additionally, it highlights the distinct cell types where these accessible regions have been recognized as the human-gained (presented on the right). **(E)** Distribution ratio of genomic annotations for various classes of accessible regions (ARs) identified in RG-neu and IPC. **(F)** Distribution of cell type conservation score for CARs, PGARs and HGARs in the IP cells. **(G)** Bar plot illustrates the percentage overlap of ARs from different categories with the human-gained enhancers (HGE) sourced from (*38*). **(H)** Heatmap illustrates the evolutionary origins of the RG-neu HGARs. **(I)** Bar plot illustrates the distribution of evolutionary origin categories for HGARs detected in different cell types. **(J)** Enrichment of the Tn5 insertion signals in humans, macaques, and mice around the cis-regulatory elements (CREs) predicted to be linked to the conserved marker genes, the primate-specific marker genes, and the human-specific marker genes.

**Fig. S13.**
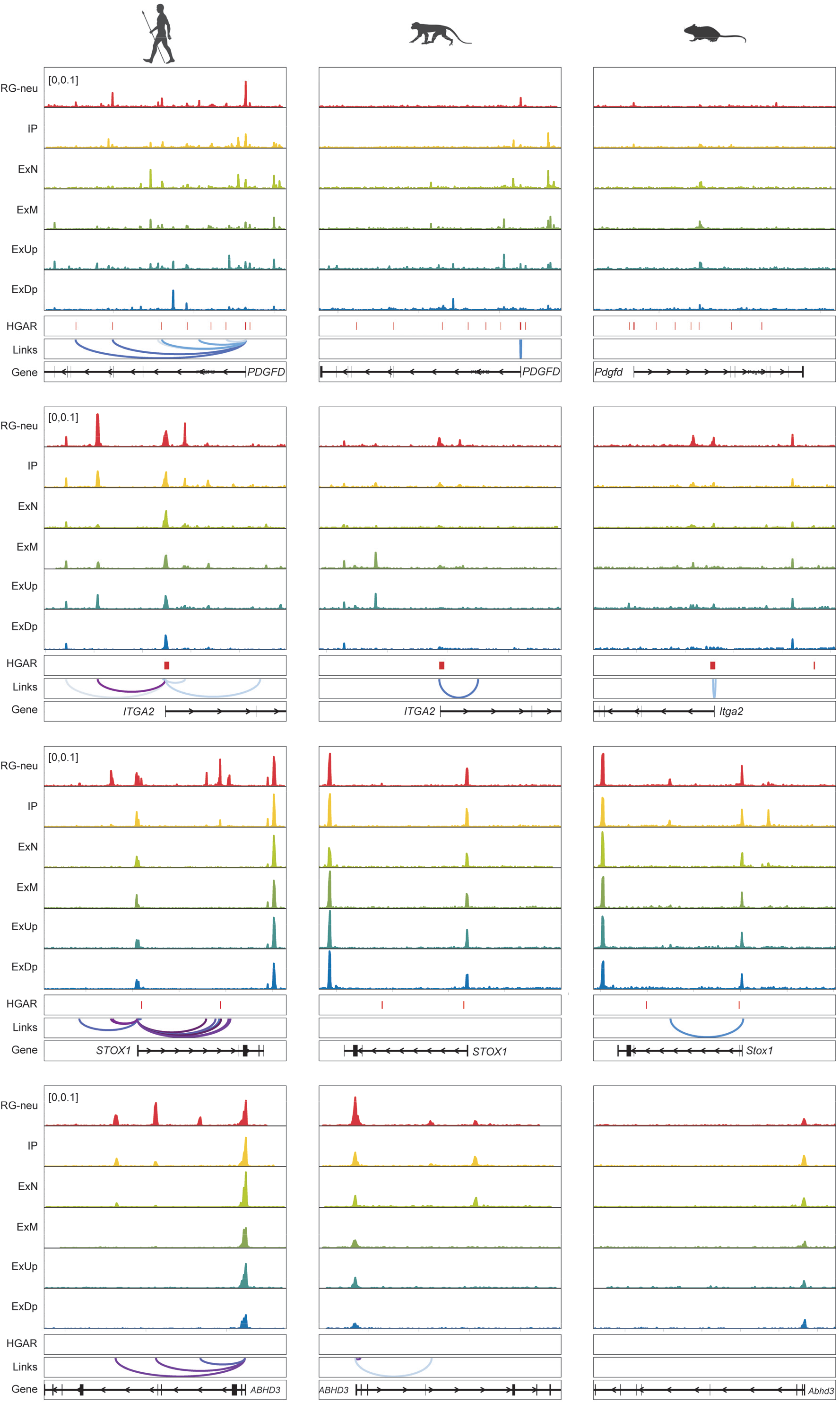
Regulatory mechanism underlying the human-specific RG-neu markers. Chromatin accessibility landscape and the predicted peak-to-gene links surrounding the human-specific RG-neu marker genes.

**Fig. S14.**
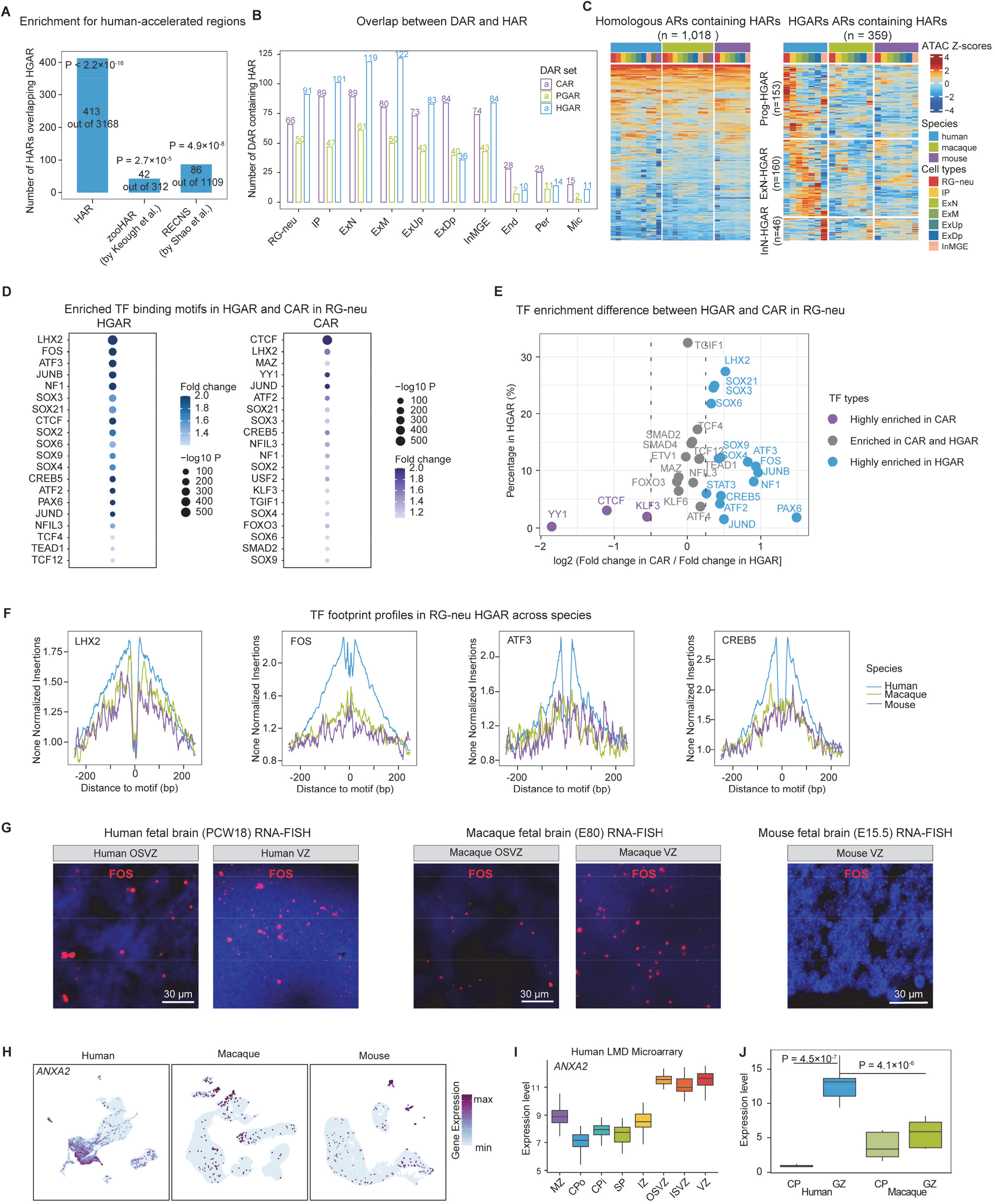
Genetic basis underlying the evolutionary innovations of the human cortex. **(A)** Bar plot illustrates the number of HARs from different studies overlapped with HGARs. **(A)** The numbers of HGARs, PGARs and CARs overlapped with HARs across different cell types. **(A)** Heatmap shows the chromatin accessibility score of the homologous accessible regions (left) and HGARs (right) containing HARs across different cell types among the three species. **(A)** TF binding motif enrichment of HGARs and CARs detected in RG-neu. **(B)** TF binding motif enrichment difference between HGARs and CARs detected in RG-neu. **(C)** TF footprint profiles of the HGAR-enriched TFs in RG-neu. **(D)** RNA FISH shows FOS expression in the fetal cortex across species. **(E)** UMAP plot illustrates the human-specific expression pattern of *ANXA2* in RG-neu. **(F)** and **(J)** *ANXA2* expression in different laminae of the human and macaque fetal cortex.

## Notes

### Competing Interest Statement

The authors have declared no competing interest.

